# Clusternomics: Integrative Context-Dependent Clustering for Heterogeneous Datasets

**DOI:** 10.1101/139071

**Authors:** Evelina Gabasova, John Reid, Lorenz Wernisch

## Abstract

Integrative clustering is used to identify groups of samples by jointly analysing multiple datasets describing the same set of biological samples, such as gene expression, copy number, methylation etc. Most existing algorithms for integrative clustering assume that there is a shared consistent set of clusters across all datasets, and most of the data samples follow this structure. However in practice, the structure across heterogeneous datasets can be more varied, with clusters being joined in some datasets and separated in others.

In this paper, we present a probabilistic clustering method to identify groups across datasets that do not share the same cluster structure. The proposed algorithm, Clusternomics, identifies groups of samples that share their global behaviour across heterogeneous datasets. The algorithm models clusters on the level of individual datasets, while also extracting global structure that arises from the local cluster assignments. Clusters on both the local and the global level are modelled using a hierarchical Dirichlet mixture model to identify structure on both levels.

We evaluated the model both on simulated and on real-world datasets. The simulated data exemplifies datasets with varying degrees of common structure. In such a setting Clusternomics outperforms existing algorithms for integrative and consensus clustering. In a real-world application, we used the algorithm for cancer subtyping, identifying subtypes of cancer from heterogeneous datasets. We applied the algorithm to TCGA breast cancer dataset, integrating gene expression, miRNA expression, DNA methylation and proteomics. The algorithm extracted clinically meaningful clusters with significantly different survival probabilities. We also evaluated the algorithm on lung and kidney cancer TCGA datasets with high dimensionality, again showing clinically significant results and scalability of the algorithm.

**Author Summary:** Integrative clustering is the task of identifying groups of samples by combining information from several datasets. An example of this task is cancer subtyping, where we cluster tumour samples based on several datasets, such as gene expression, proteomics and others. Most existing algorithms assume that all such datasets share a similar cluster structure, with samples outside these clusters treated as noise. The structure can, however, be much more heterogeneous: some meaningful clusters may appear only in some datasets.

In the paper, we introduce the Clusternomics algorithm that identifies groups of samples across heterogeneous datasets. It models both cluster structure of individual datasets, and the global structure that appears as a combination of local structures. The algorithm uses probabilistic modelling to identify the groups and share information across the local and global levels. We evaluated the algorithm on both simulated and real world datasets, where the algorithm found clinically significant clusters with different survival outcomes.

## Introduction

Current growth in high-throughput analysis methods in bioinformatics gives access to many different types of data measuring behaviour in complex biological systems. It is now possible to observe multiple levels of a complex biological process simultaneously. For example, we can look at DNA copy number changes, gene expression patterns, epigenetic features such as DNA methylation, and protein expression - all giving a view of different aspects of the underlying process. Analysis of such data is non-trivial because of the need to integrate information from the different levels. Integrative methods that look at the combined effects of several levels of biological processes in the cell have the potential to identify more features that lead to complex phenotypes.

We introduce a novel algorithm for integrative clustering for heterogeneous multilevel data: context-dependent clustering. In general, integrative clustering is the task of identifying clusters in a set of related datasets across the same samples.

As a motivating example, we use the problem of cancer subtyping. Cancer is a heterogeneous process where even cancers originating from the same tissue behave differently in relation to their original driver mutations [1]. Cancer subtyping is the task of identifying types of cancer which are typically associated with different patient outcomes, therapy responses or tumour aggressiveness. Finding such cancer subtypes then allows identification of the differences between their molecular behaviour. By integrating different levels of data we can get a better understanding of the interplay of different steps in cancer biochemical pathways.

In the cancer subtyping problem, we use integrative clustering to characterise different types of cancer, based on the tumours’ genomic and other omic profiles. The tumour samples are simultaneously analysed using different technologies, for example gene expression, methylation, and sequencing, yielding a set of related datasets. Integrative clustering looks for a partitioning of tumour samples based on their exhibited behaviour across the datasets.

Compared to standard clustering methods, the main challenges in integrative clustering come from the fact that individual datasets are not directly comparable: each of them describes a different aspect of the underlying biological process. Datasets may even have different data types - continuous gene expression values, binary presence/absence of chromatin modifications or even time-series observations and clinical markers.

Because each data set originates from a different context, clusters obtained from different datasets are not consistent in general. Cluster membership and/or the number of clusters may differ between datasets. This discrepancy originates both from noise and from biological heterogeneity. For example, Ovaska et al. [2] report unexpectedly poor concordance between gene amplification, expression of the genes from the amplicons, and patient survival in a study on childhood brain tumour glioblastoma multiforme. Kristensen et al. [3] analysed results from dataset-specific clustering analyses on different types of omic data and note that only a minority of the samples were grouped together across all datasets.

These examples show that the notion of a cluster itself is ambiguous in the context of integrative clustering. Most existing methods aim to extract a common cluster structure that is shared across all datasets. However, due to the multiple different sources of heterogeneity, biases and high level of noise, a single cluster structure that would be common to all datasets may not exist or it may not be identifiable in some of the datasets. The assumption of a common cluster structure is well suited to model only the processes that show consistent behaviours across all datasets. However, this assumption is limiting in modelling complex heterogeneous processes such as cancer.

Using a simple example from breast cancer research, both increase in copy number of the HER2 gene, and increase in expression of the corresponding protein are correlated with poor prognosis in patients [1]. One would naively expect that the oncogene copy number amplification leads to over-expression of its protein. However, the relation is not always this straightforward: some tumours over-express HER2 even without the original gene amplification [1]. The over-expression is apparently achieved through a mechanism other than a change in gene copy number. However, the samples would not be distinguishable based on gene expression data alone, but can be separated by taking genomic information into account as well.

By assuming a single common cluster structure, these heterogeneous situations are difficult to model. A set of samples may systematically belong to two distinct clusters in one dataset (copy number aberration) and to a single joint cluster in a second dataset (mRNA gene expression). This example shows that real biological processes display different behaviours in different contexts, which leads to different cluster arrangements.

The existing algorithms for integrative clustering assume that there is only a single common cluster structure across all analysed datasets. Individual samples then either follow this structure or they are considered to be a noise. This treatment of inconsistent samples is suitable for technical noise which does not bear biological significance, but it can miss cases where the difference is biologically meaningful.

The proposed context-dependent clustering model assumes instead that a single biological process (such as cancer) looks and behaves differently in different *contexts*. A context might represent different stages of the process (early versus late carcinogenesis), or different data domains (DNA modifications, mRNA expression, etc.).

### Existing methods assume a common structure

Here we look at some of the existing approaches to integrative clustering and how they use the assumption of a common clustering structure.

The iCluster algorithm [4, 5] uses a Gaussian latent variable model to infer clusters. It assumes that there is a common set of latent cluster membership variables across all datasets. Differences in structure between different datasets are accounted for only via individual noise terms, which correspond to within-dataset variances. iCluster uses the *k*-means algorithm to extract the actual cluster assignments given latent variable values.

Using this approach, it is possible to obtain clusters where the data are well separated across all datasets. However, if two clusters are joined together in one dataset and separate in another dataset, this leads to ambiguous latent variable values and iCluster fails to infer the correct partitioning. Also, using *k*-means for cluster inference from latent variable values makes the algorithm strongly dependent on the correct specification of the number of clusters. We demonstrate this behaviour on a simulated dataset later in the paper.

A similar algorithm to iCluster is the Joint and Individual Clustering (JIC) [6]. This algorithm uses the connection between principal component analysis and *k*-means clustering to integrate data from multiple datasets and infer both joint clusters and dataset-specific clusters. The dataset-specific clusters are estimated so that they are independent of the joint clusters, and thus capture independent structure that exists only within individual datasets. This algorithm still uses the assumption of one global cluster structure across the datasets, which is augmented by additional cluster structures within individual datasets that are independent of the overall global structure.

Bayesian consensus clustering (BCC) [7] relaxes the assumption of a common cluster structure by allowing different datasets to follow individual *local* clustering models. Apart from the local clusters, BCC introduces a *global* cluster structure as well. This depends on the local clusters through a parameter *α*, which regulates how much the local clusters correspond to the global clusters. The split between global and local structure brings flexibility in modelling inconsistent cluster structures across many datasets. The consensus global cluster assignments then define a common clustering based on partial agreement between context datasets.

However, this again assumes that there is a common set of clusters which are exhibited in a similar way across all datasets. Deviances from this structure are considered to be dataset-specific noise, which might lead to omission of any additional structure that is present in the data.

Multiple Dataset Integration (MDI) [8] uses a different approach. This model does not assume a common clustering but it looks only at pair-wise relations between datasets. In this formulation, every dataset can have a flexible set of clusters, and a pairs of samples are considered *fused*, if they belong to the same set of pairs of clusters across datasets. This limits the interpretability of a solution. Although the model allows enough flexibility for combinations of clusters, it does not explicitly encourage any sharing of clusters across more than pairs of datasets.

Another approach is taken by the Similarity Network Fusion (SNF) [9]. This algorithm uses an iterative approach on dataset-specific similarity networks that intensifies strong similarities and diminishes weak similarities across samples. This again implies the existence of a common structure that is consistent across the datasets.

### Context-dependent integrative clustering

We introduce the context-dependent clustering model to address the issues outlined in the introduction. Compared to existing integrative clustering algorithms, the proposed model does not generally assume a single partitioning of the data that is consistent across heterogeneous datasets.

If a process is allowed to follow different cluster structures across several contexts, we have to modify our notion of clustering. We assume that there is a set of clusters within each context (dataset), but these clusters do not generally correspond directly to each other. However, some dependency between the clusters is expected:

1. Clustering structure in one context should influence clustering in other contexts. If two samples are clustered together in one context, they should be more likely to be clustered together in other contexts as well.
2. Different degrees of dependence should be allowed between clusters across contexts. For example, datasets may not share the same numbers of clusters. We should be able to model cases where all contexts share the same cluster structure, as well as cases where the clusters are completely independent in the different contexts.

Using the example of the HER2 oncogene from the Introduction, there are two levels of complexity we can analyse. If we consider only tumour samples which over-express HER2, the first level is formed by the individual contexts: there are two clusters in DNA amplification dataset (amplified or neutral), and one cluster when we consider mRNA expression of HER2 (over-expressed). If we assumed a globally consistent cluster structure, we would have to either artificially split the cluster of samples which over-express HER2, or artificially join the two clusters with different copy number aberrations. When we look at the problem from the *global* level, we get two groups of samples that behave in a different way across contexts. The behaviours are defined by different *combinations* of context-specific clusters. The concepts are illustrated in Fig. 1.

**Figure 1.**
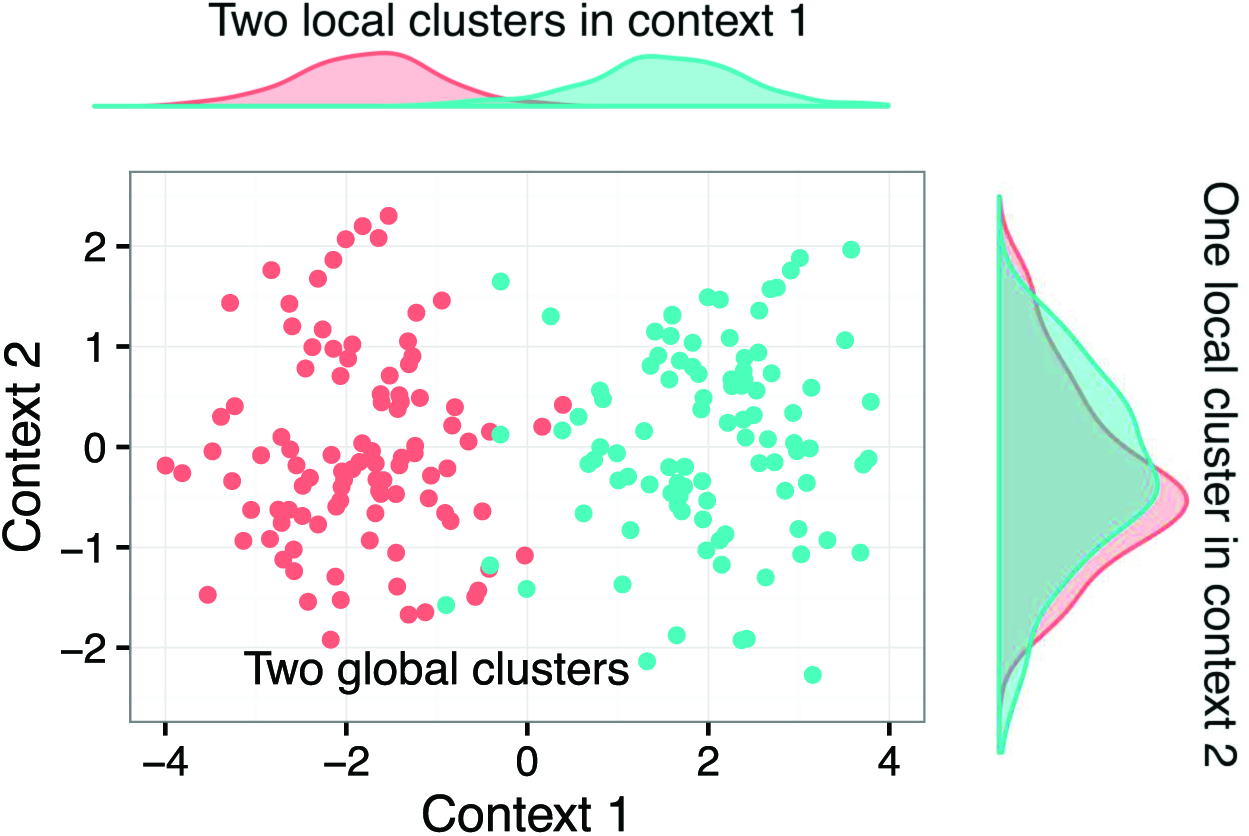
Illustration of heterogeneous cluster structures in two contexts (datasets). Each context corresponds to different data source (gene expression, ribosome profiling, proteomics etc.) describing the same set of biological samples. In the first context, there are two distinct clusters on the local level. In the second context, there is only a single local cluster. From the overall perspective, there are two global clusters defined by the combined behaviour across the two contexts.

The context-dependent clustering model as proposed here uses a Bayesian clustering framework to infer both the local structure within each dataset, as well as the global structure which arises from the combination of cluster assignments. We use probabilistic framework, with hierarchical Dirichlet mixture models to model clusters across the contexts. We fit the model using Gibbs sampling. Details of the model formulation and inference algorithms are presented in the Methods section.

In the next section we focus on a comparison of the proposed algorithm with alternative context dependent clustering approaches on simulated as well as on real-world datasets.

## Results

We evaluate the proposed context-dependent clustering model both on simulated dataset to compare its performance with other currently used methods, and on real world datasets. The simulated example demonstrates that the proposed model identifies clustering structure within datasets where the dependence of individual contexts varies. The applications to the real world datasets (breast cancer, lung cancer and kidney cancer) from The Cancer Genome Atlas (TCGA) show that the algorithm identifies biologically meaningful structure, which is characterized by significantly different survival outcomes. Finally, we demonstrate the robustness of the results to changes in parameter settings.

### Case Study: Simulated Datasets with Heterogeneous Structure

In this section, we look at how the proposed context-dependent clustering method performs in a situation where there is no single common clustering structure. We compare its performance with popular alternative clustering algorithms on a set of simulated data with various degrees of dependence between contexts.

The simulated data are formed by two 1-dimensional datasets, which represent a dataset with two contexts. Each context contains two clusters: both clusters are normally distributed with unit variance but with different means. Cluster 1 in both contexts is centred at −2 and Cluster 2 is centred at 2. The two clusters are well separated and they do not overlap significantly. We combine the clusters to simulate different degrees of independence between context-specific cluster structures.

We generate 100 data sets each with 200 samples according to the following procedure. For each data set we sample a mixture probability *p* uniformly from the interval (0, 0.5). For the first group of 100 samples we sample each of the two context values independently from 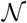(−2, 1) or from 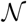 (2, 1) with probability 1 − *p* and *p*, respectively, that is, since *p* < 0.5 mostly from Cluster 1, but occasionally also from Cluster 2. For the next group of 100 samples we sample the each of two context values independently from 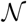(−2, 1) or from 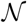(2, 1) with probability *p* and 1 − *p*, that is, mostly from Cluster 2 this time, but occasionally from Cluster 1.

For mixture probabilities *p* close to 0, the clusterings of the two contexts largely agree by putting the first 100 samples into Cluster 1, and the rest into Cluster 2, leading to two global clusters. At the other extreme, if *p* is close to 0.5, cluster assignments are more or less random and there is little agreement between contexts, leading to four global clusters.

Fig. 2 shows data sampled from the two opposing scenarios. If the data distributions are fully dependent across the contexts, we get two global clusters (Fig. 2a). Because the data are assigned to the same cluster in both contexts, there is a single common cluster structure. On the other hand, if the two contexts are independent, we get four global clusters (Fig. 2b) which correspond to the combinations of context-specific cluster assignments. This scenario is difficult to model if we look at individual contexts separately.

**Figure 2.**
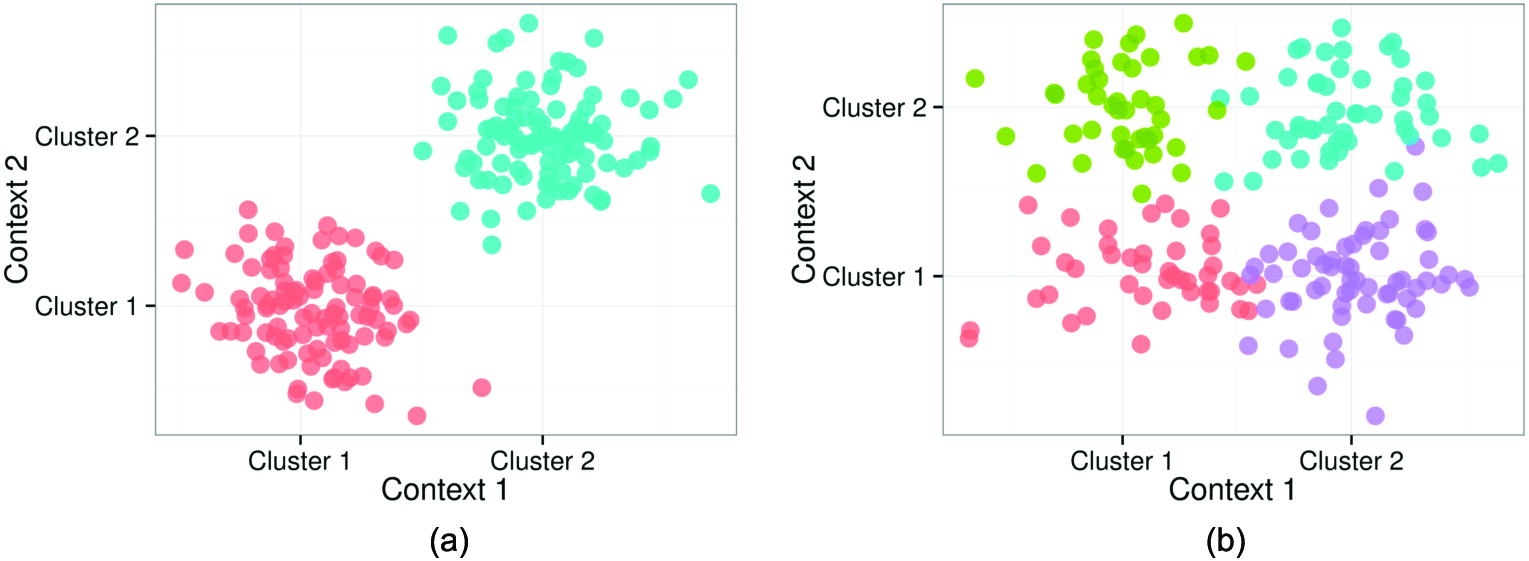
Example of the simulated data for *p* = 0 and *p* = 0.5, which show different degrees of dependence. The *x* axis corresponds to the data in the first dataset (context), the *y* axis represents the data in the second dataset (context). The two subfigures show the two extreme situations: (a) For *p* = 0, we get two global clusters. Cluster membership is fully dependent on each other in both datasets. (b) For *p* = 0.5, we get four global clusters, where cluster membership in one dataset is fully independent on cluster membership in the second dataset.

#### Results

We use the simulated data to evaluate the following integrative clustering algorithms:

1. The proposed Context-Dependent Clustering (Clusternomics)
2. Bayesian Consensus Clustering (BCC) [7]
3. Multiple Dataset Integration (MDI) [8]
4. iCluster [4, 5]
5. Similarity Network Fusion [9]

Methods 1, 2 and 3 are Bayesian probabilistic methods fitted using MCMC sampling, 4 uses a latent variable model fitted with an EM algorithm combined with *k*-means clustering and finally 5 is an algorithm based on summarised distances between samples.

We ran the listed methods on the 100 simulated datasets. Details on the settings of the individual algorithms are given in S1 Appendix. The results are only comparable on the level of global clustering due to the properties of the algorithms themselves: SNF and iCluster only compute the global clustering across all contexts. BCC and MDI have a notion of local clustering within each context, but they do not allow the number of clusters to differ between the global and local level. All methods except for MDI use the number of global clusters as their input: we set this equal to 4.

We use the adjusted Rand Index (ARI) [10] to compare the true cluster assignments used to generate the data, and the results estimated by each algorithm. The ARI measures agreement between two partitions of the same set of samples; the value of 1 corresponds to complete agreement between two cluster assignments, and 0 means the agreement between partitions is caused by chance. Fig. 3 shows ARI values comparing the different algorithms for the 100 simulated datasets.

**Figure 3.**
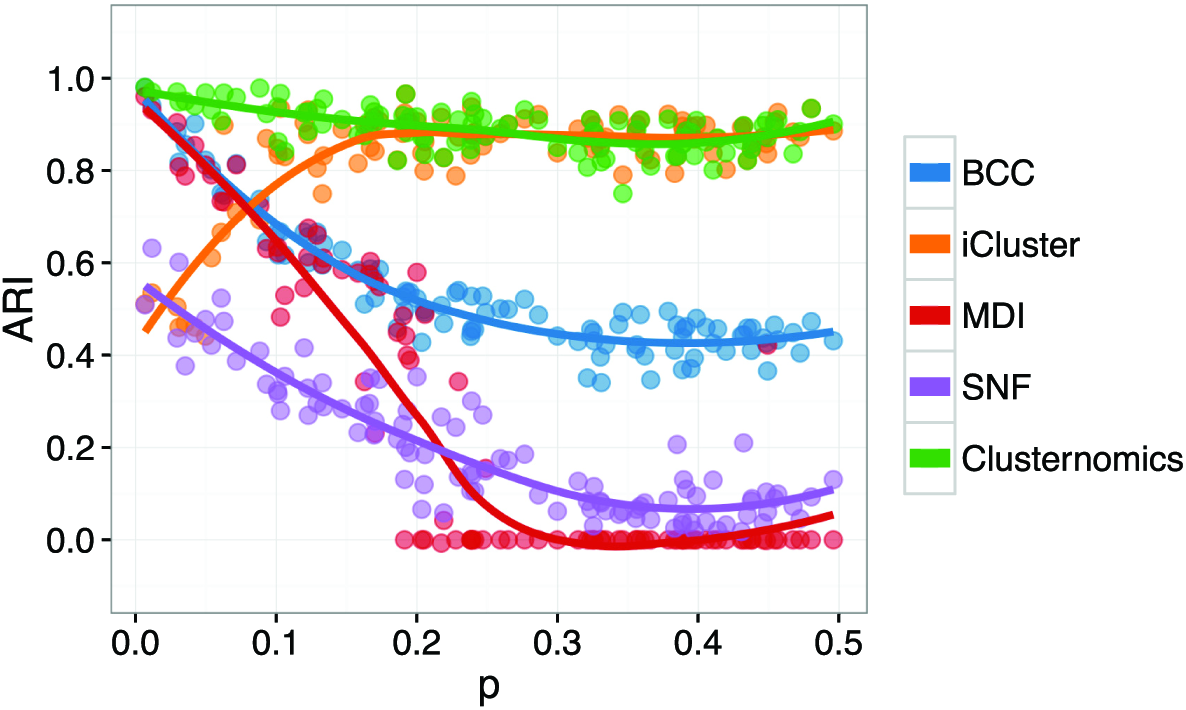
ARI comparing global clustering of simulated datasets for varying values of *p* (see Fig. 2). Each point corresponds to the corresponding algorithm applied to one dataset, the plot shows also the loess curve for each method. Higher values correspond to better agreement between the estimated cluster assignments and the true cluster membership.

For small values of *p*, which correspond to two fully dependent global clusters, all probabilistic methods Clusternomics, BCC and MDI perform similarly well. For higher values of *p*, which represent higher degrees of independence of clusters between contexts, Clusternomics and iCluster have the best performance. However, overall only Clusternomics is able to recover the underlying cluster structure across all different values of *p*.

The disappointing performance of MDI is caused by the algorithm allocating all the samples to a single cluster for higher values of *p*. One should note, however, that the MDI algorithm was disadvantaged in this setting because it infers the number of clusters from the data instead of using a pre-specified value.

The iCluster and SNF algorithms were also disadvantaged specifically for small values of *p* because these algorithms use k-means and spectral clustering respectively to extract the number of clusters. This makes them very sensitive to mis-specification. As an illustration, Fig. 4 shows the behaviour of all analysed algorithms when we set the number of global clusters to 5, and allowed up to 3 clusters in every context in the Clusternomics algorithm. We can see that the change in the number of clusters does not affect probabilistic algorithms but iCluster and SNF are strongly dependent on this setting.

**Figure 4.**
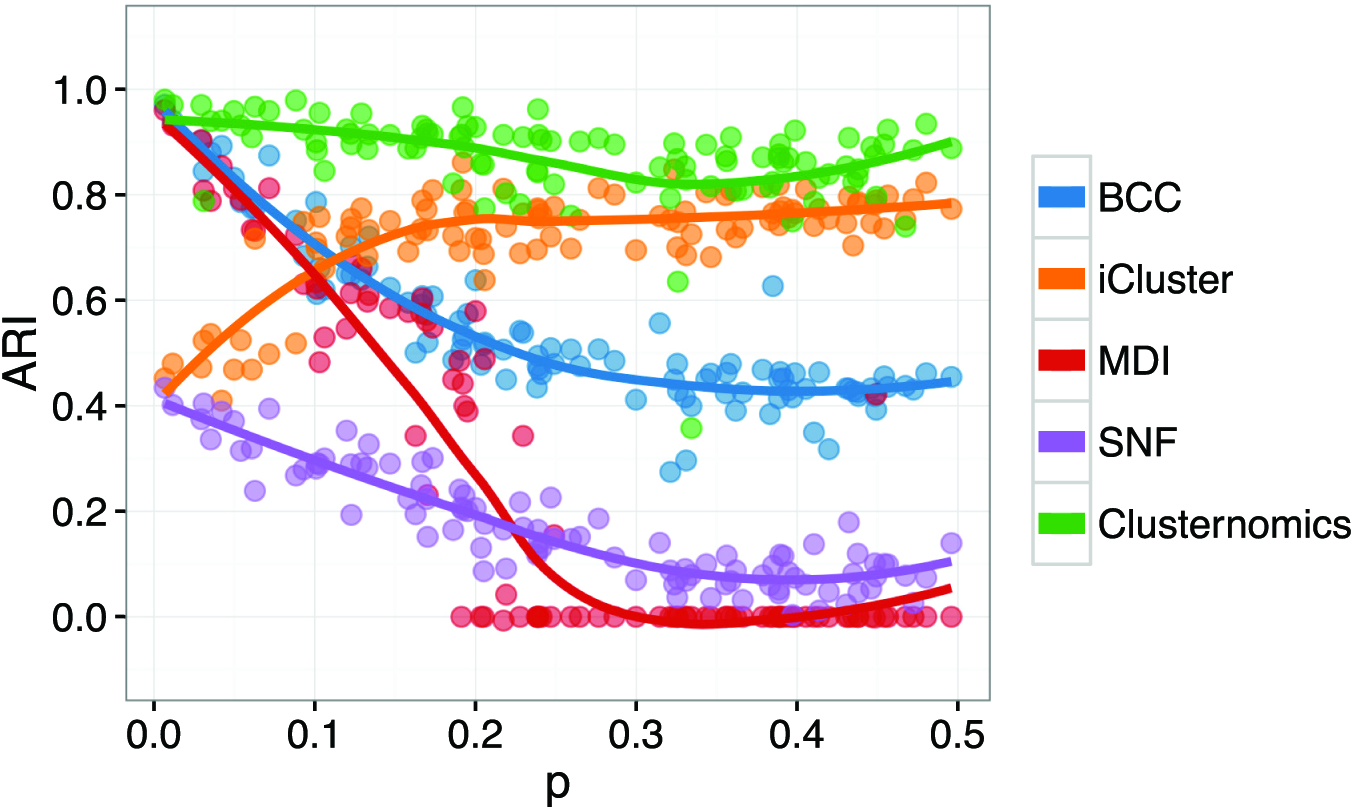
ARI comparing global clustering of simulated datasets with misspecified number of clusters for varying values of *p*, when we set the number of global clusters to 5. Each point corresponds to a corresponding algorithm applied to one dataset, the plot shows also the loess curve for each method. Higher values correspond to better agreement between the estimated cluster assignments and the true cluster membership.

In real world applications, it is often very difficult to estimate the number of clusters correctly. Although Dirichlet process-type methods are inconsistent in terms of the true number of clusters [11], their results are generally consistent in the number of large clusters that appear in the data.

We also provide an additional comparison in Sl Appendix with an ad-hoc integrative clustering, where we first cluster data in each context individually, and then we construct *global* clusters manually as a combination of *local* cluster assignments. For the small simulated dataset, the results of this ad-hoc integration are equivalent to Clusternomics. They differ in larger real-world applications, where the probabilistic integrative model encourages data points from different contexts to share global clusters, as opposed to crude manual construction of global clusters.

### Case Study: Discovering Subtypes in Breast Cancer

In this section we apply the context-dependent clustering to a breast cancer dataset obtained from The Cancer Genome Atlas (TCGA) database. The dataset contains measurements of 348 patients diagnosed with breast cancer, and it comprises of four different data types: DNA methylation for 574 probes, RNA gene expression for 645 genes, expression for 423 microRNA molecules and reverse-phase protein array (RPPA) measurements for 171 proteins.

The dataset was originally presented in Koboldt et al. [12], and it was also previously used to evaluate integrative clustering methods in Lock and Dunson [7]. An advantage of the dataset is that it also contains clinical information on the patients including their survival. This information can be used to validate the clustering results.

We used the proposed context-dependent clustering algorithm to identify subtypes of breast cancer across the four data contexts. We applied the proposed algorithm with several different settings of both the *global* number of clusters, and the *local* context-specific number of clusters. The details of the algorithm's settings are provided in S1 Appendix.

To provide an overview of the results, we first look at a specific clustering obtained from the model to illustrate the working of the algorithm. Then we look at consistency of the results with respect to varying number of clusters, and also at results from two other cancer datasets from TCGA.

To illustrate the results from the model, we use an example with the number of local clusters set to 3 in all the four contexts (gene expression, DNA methylation, miRNA expression and RPPA) and 18 global clusters. The number of global clusters acts as an upper bound on the number of clusters that can be represented in the data by the clustering model. The number of clusters that the model identified in the presented result was 16. Fig. 5 shows the size distribution of these 16 clusters. There are several larger clusters and a larger number of smaller clusters.

**Figure 5.**
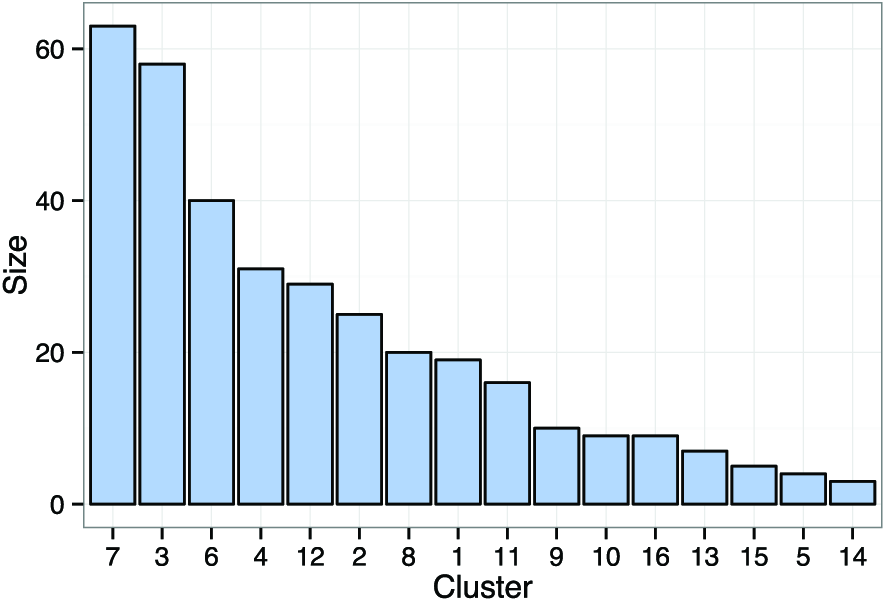
Sizes of global clusters identified in the breast cancer dataset from TCGA, using the model with 3 context-specific clusters and up to 18 global clusters.

The global combinatorial clusters identified by the model showed clinical significance in terms of survival probabilities. Fig. 6 shows the different survival curves corresponding to each of the clusters. The survival probabilities in each cluster are significantly different with *p* = 0.038 using the log-rank test with a null hypothesis that assumes that the survival rates are the same. Here, identification of the global structure helps clustering within individual datasets. Looking at the local cluster structure within each dataset, the differences in cluster survival curves are not individually significant. By looking at the overall global structure, we can identify groups that are characterized by different survival outcomes.

**Figure 6.**
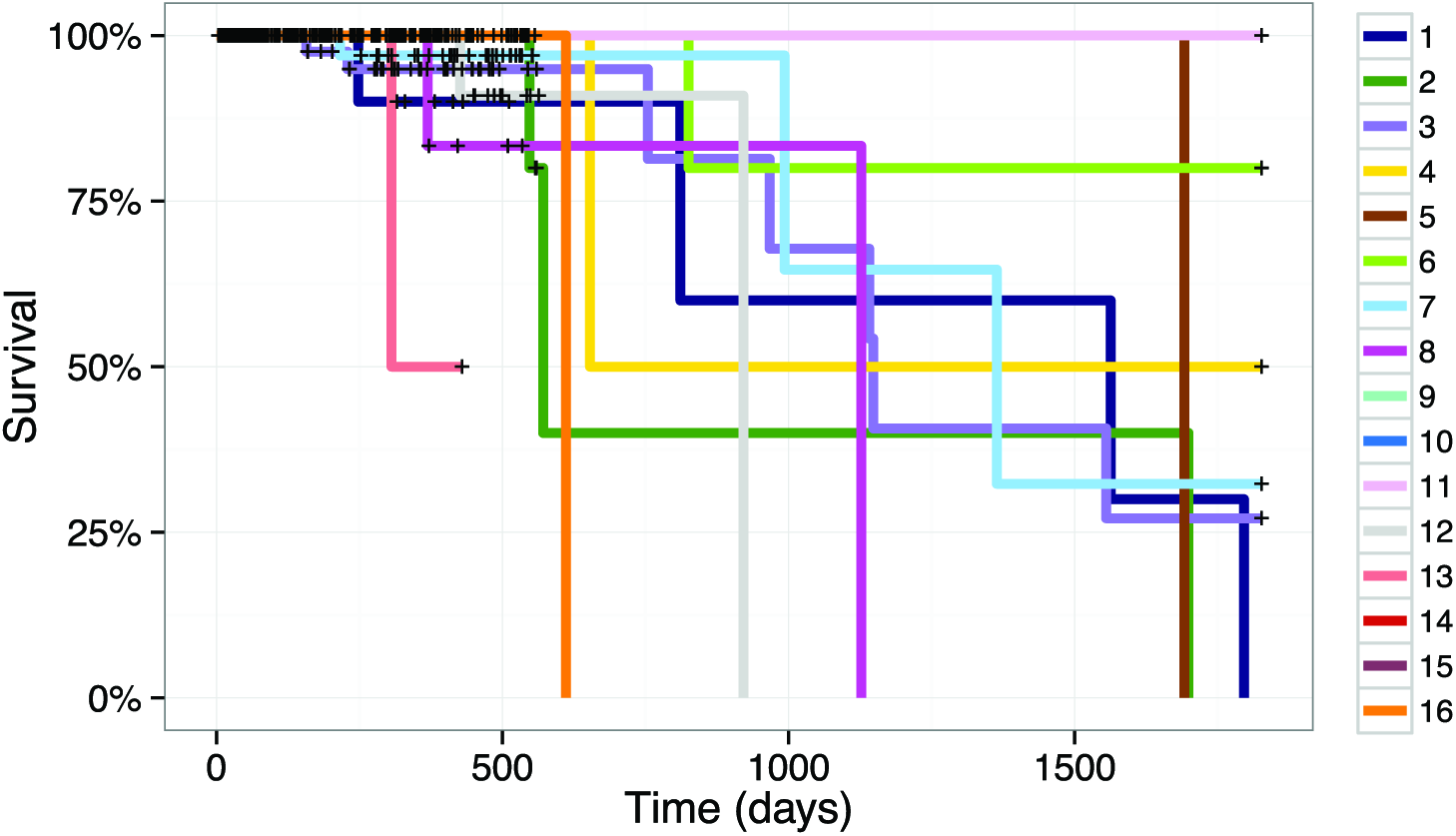
Survival curves for global clusters in the breast cancer dataset from TCGA, using the model with 3 context-specific clusters and up to 18 global clusters. The differences between the survival curves are significant with *p* = 0.0382 using the log-rank test.

Fig. 7 shows the global clusters as they appear in the four individual contexts in the first two principal components. The clusters are better defined for the gene expression and the protein assay context. However, the samples corresponding to each cluster also occupy distinguishable regions in the remaining contexts. Because the plots show only the first two principal components, we also provide an over view of variance captured by these components in S1 Appendix.

**Figure 7.**
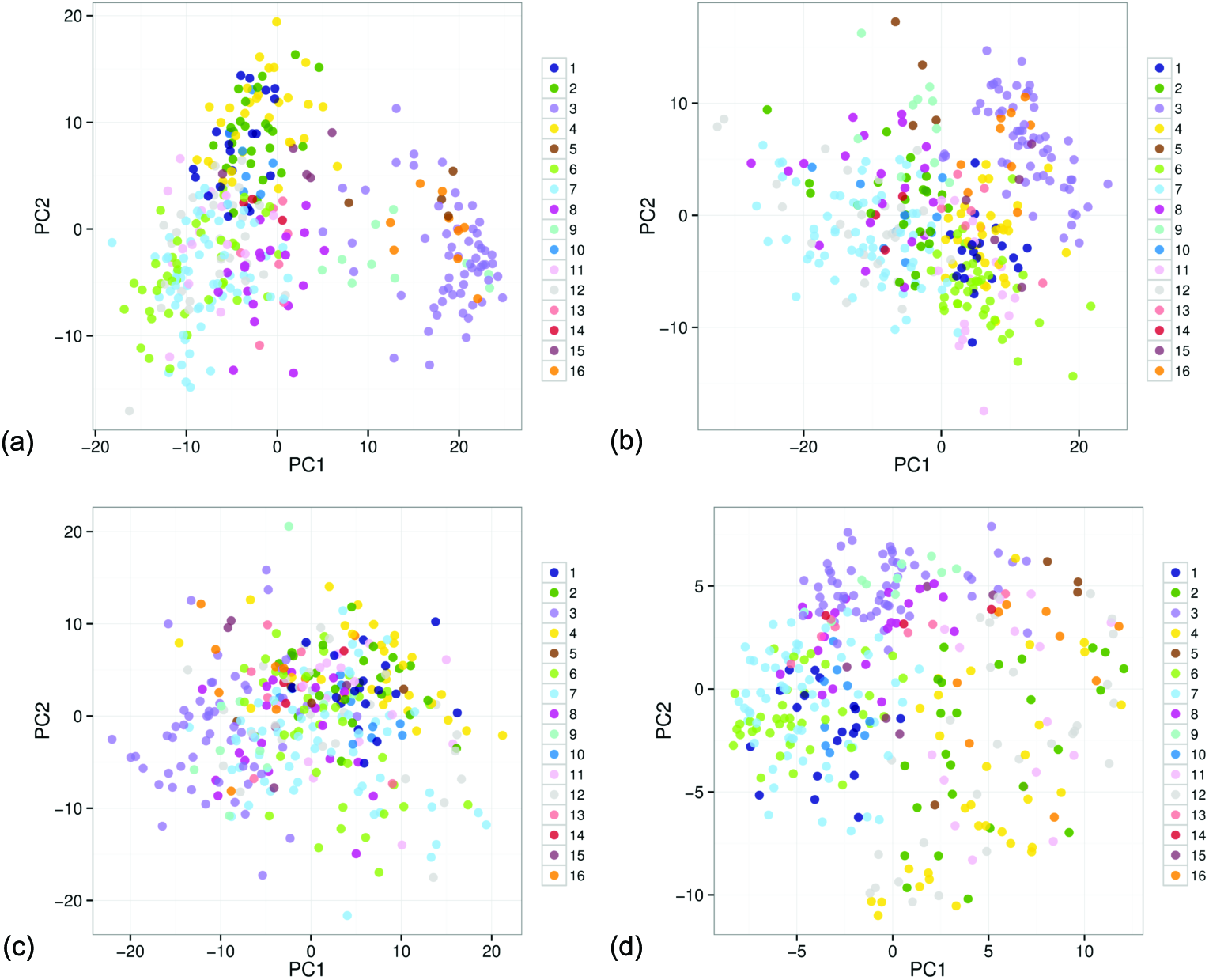
PCA projection of the global clusters in individual contexts in the breast cancer dataset from TCGA, from the model with 3 context-specific clusters and up to 18 global clusters. The colours correspond to the colours used in the survival curves in Fig. 6. (a) Gene expression context. (b) DNA methylation context. (c) miRNA expression context. (d) RPPA context.

In contrast to the global clusters in Fig. 7, Fig. 8 shows the inferred local clusters within each dataset. The clusters are well defined in all contexts except the miRNA expression dataset, where the model did not identify any structure. However, by sharing information from other contexts we see that some of the global clusters are distinguishable in this context as well. For example, in Fig. 7c Cluster 3 (violet) occupies a distinguishable region in the miRNA context although the dataset itself does not show any pronounced clustering tendency within the first two principal components. This indicates that the identified clusters represent some feature of the underlying process even within datasets that, by themselves, contain only a vague structure.

**Figure 8.**
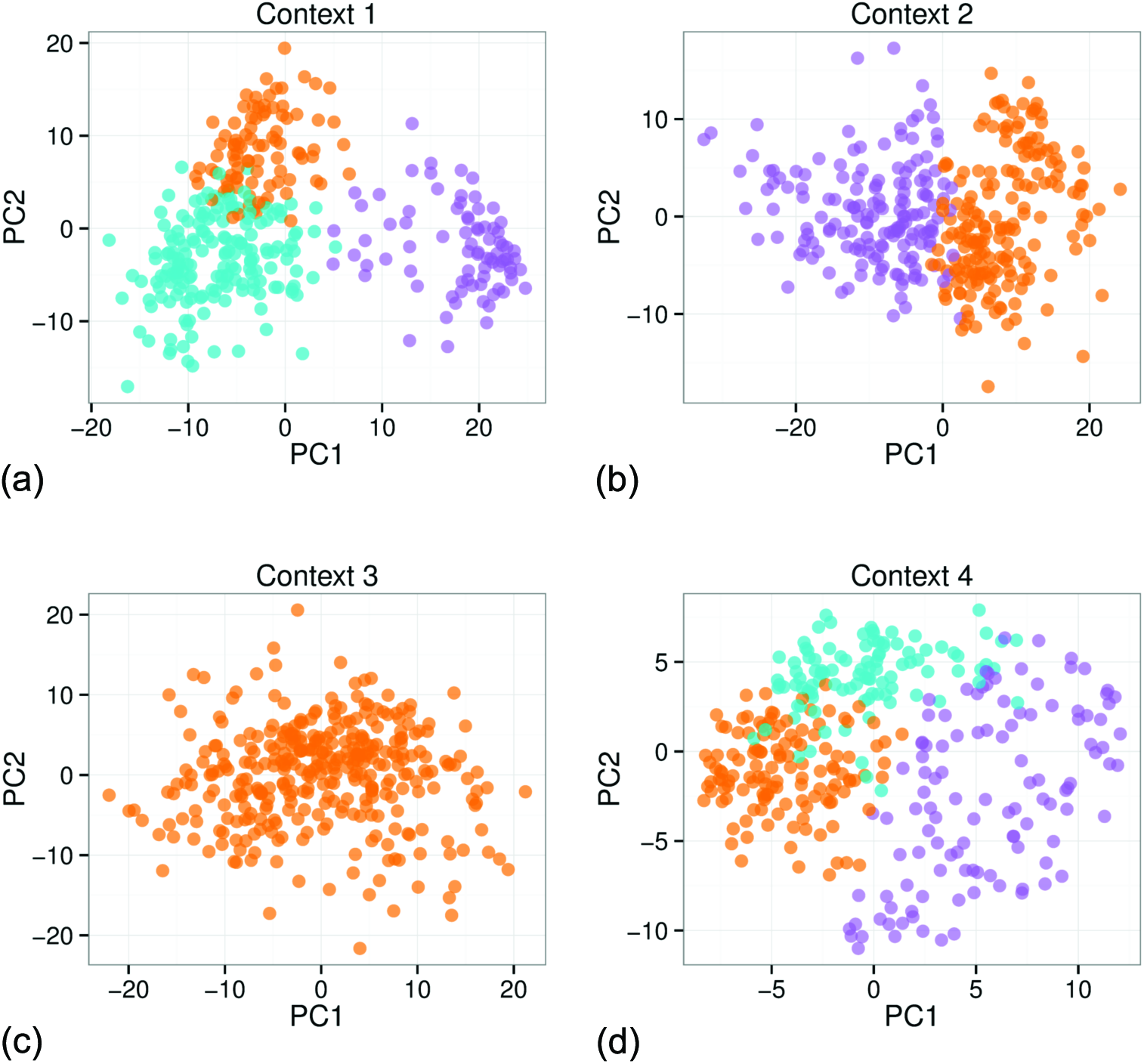
PCA projection of the local clusters identified in individual contexts in the breast cancer dataset from TCGA, from the model with 3 context-specific clusters and up to 18 global clusters. (a) Context 1 represents the gene expression dataset which contains three local clusters. (b) Context 2 represents the DNA methylation dataset and contains two local clusters. (c) Context 3 represents the miRNA expression dataset with only 1 cluster. (d) Context 4 corresponds to the RPPA dataset, which contains three local clusters.

To highlight some of the features of the presented context-dependent clustering model, we look at two clusters in more detail, namely clusters 1 and 6. Fig. 9 shows the two highlighted clusters. We can see that they are similar in all contexts except gene expression, where they belong to different clusters. This is the type of structure where two clusters are merged into one in some of the contexts and separate in other contexts, which we illustrated at the beginning of this chapter using the example of the HER2 gene. This type of structure relates to the interplay of different levels in the underlying biological process.

**Figure 9.**
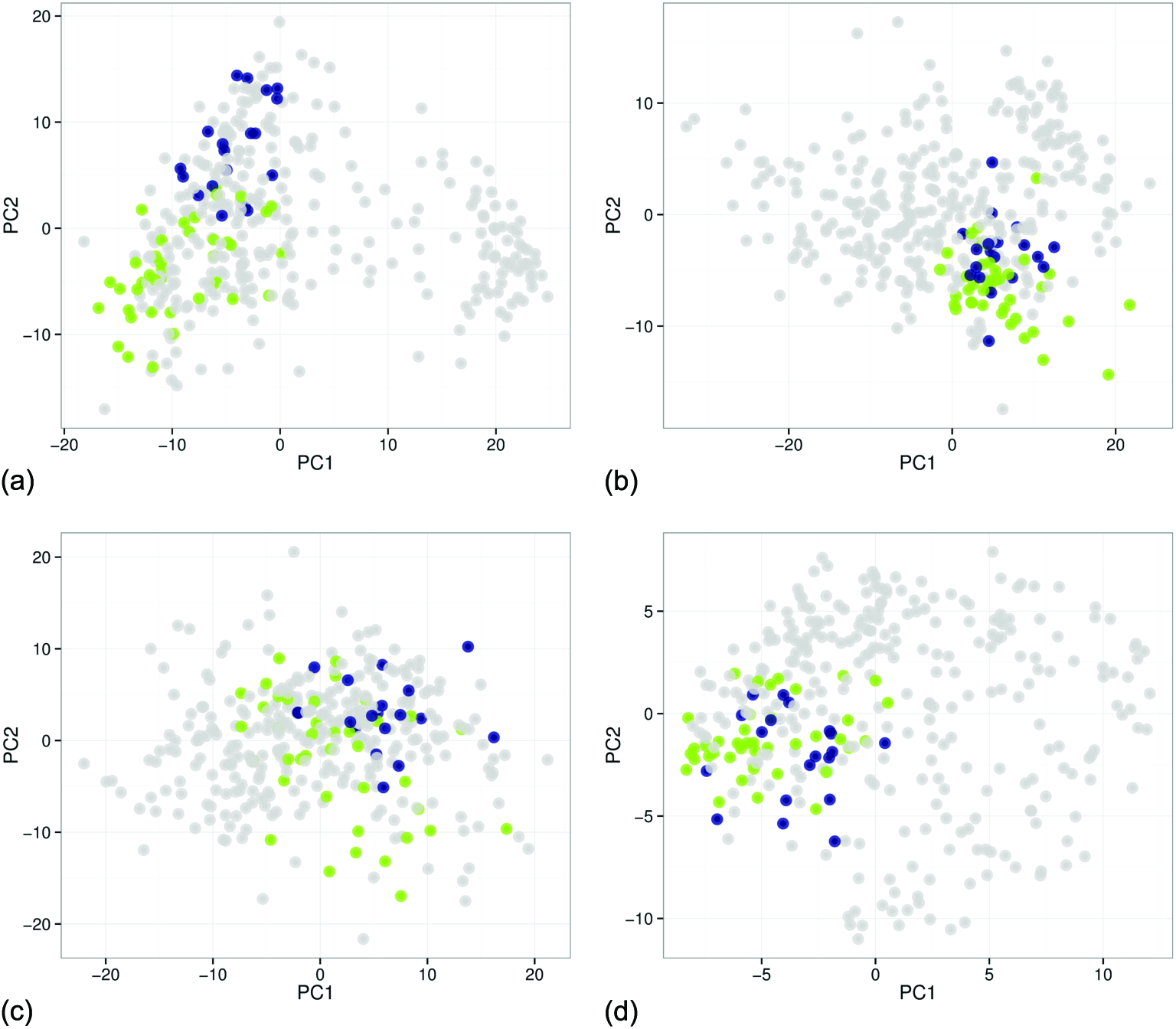
PCA projections of the global clusters identified in individual contexts in the breast cancer dataset from TCGA. The two highlighted clusters differ only in the gene expression context but they are merged in the other contexts. (a) Context 1 represents the gene expression dataset where the two clusters are separate. (b) Context 2 represents the DNA methylation dataset. (c) Context 3 represents the miRNA expression dataset. (d) Context 4 corresponds to the RPPA dataset.

The differences between the two clusters are also biologically relevant. Fig. 10 shows the highlighted survival curves for the two clusters. The green cluster (Cluster 6) contains 40 samples and has significantly different survival outcomes from the dark blue cluster (Cluster 1) containing 19 patient samples, with a log-rank test *p*-value of 0:012.

**Figure 10.**
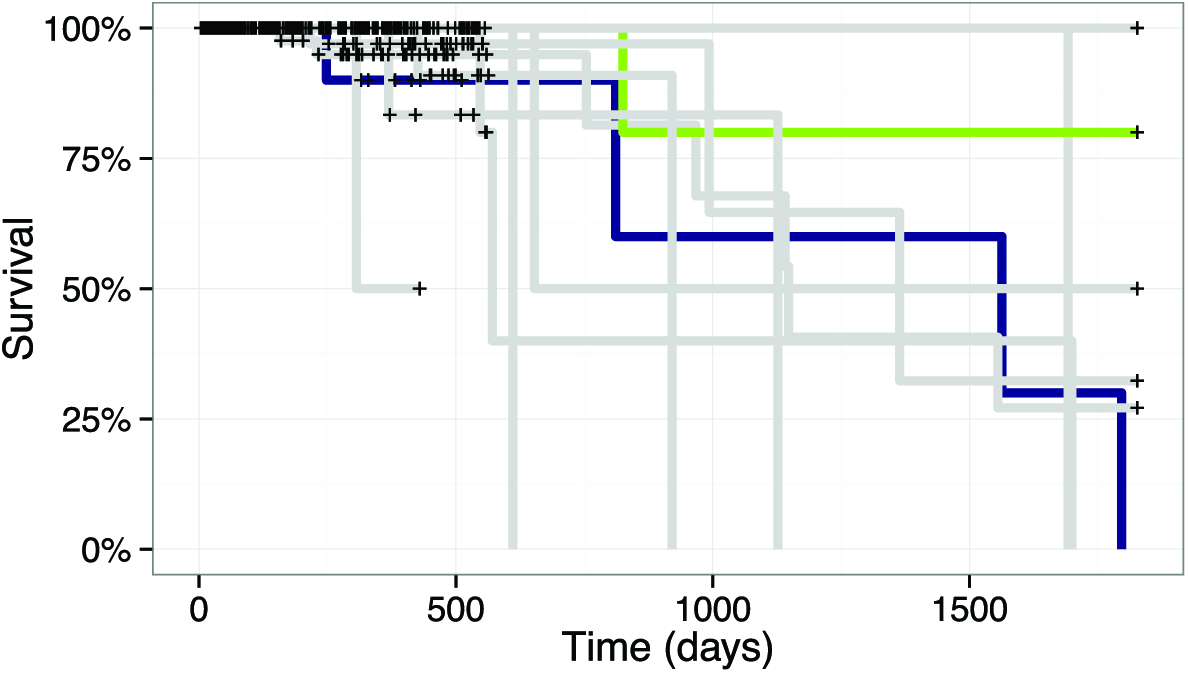
Survival curves for clusters in Fig. 9. The highlighted clusters have different survival probabilities with *p* = 0.012 under the log-rank survival model.

The presented result from the model shows that the identified structure is clinically relevant. The assumptions behind the Clusternomics algorithm also help identify combined global clusters that would be missed by other algorithms, as demonstrated on the simulated datasets.

### Stability of Inferred Clusters

The illustration of the results looked at a specific size of the model. The algorithm also allows the user to specify the number of clusters both on the local context-specific level, and on the global level. We fitted the model with varying numbers of clusters on both levels and we look at the stability of results with respect to different parameter settings. The results show that the estimated global clustering structure is stable when the number of clusters is large enough to give the model sufficient flexibility to fit the data.

Fig. 11 shows a comparison of log likelihoods of the fitted models for 10 different numbers of global clusters. The maximum of the log likelihood corresponds to the model with 18 global clusters that we highlighted as an example in the previous section. The same model also corresponds to the lowest *p*-value with respect to the differences in the survival function, as shown in Fig. 11b.

**Figure 11.**
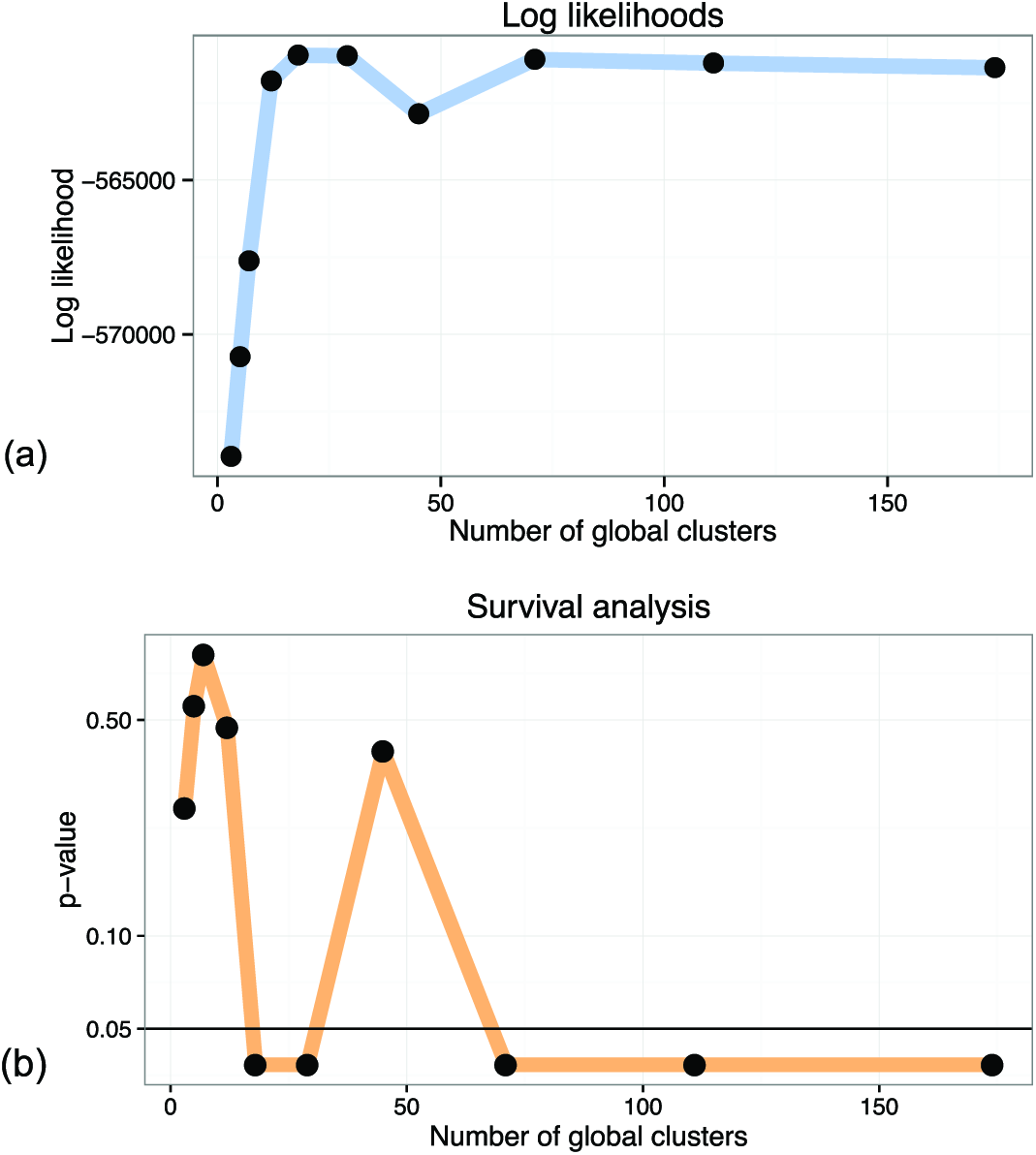
Log likelihoods (a) and survival *p*-values (b) of models with different numbers of global clusters. The first significant difference in survival corresponds to the model with the highest log likelihood.

The number of global clusters specified in the algorithm serves only as the upper limit on the number of occupied clusters. To investigate this relation we also look at the number of clusters that are actually occupied for different settings of the number of global clusters. Fig. 14 shows the posterior average number of clusters that had samples assigned to them. The figure shows both the average total number of occupied clusters across Gibbs samples for each run, and the average number of *large* clusters (clusters that have at least five samples assigned to them). The number of large clusters is more reflective of the actual underlying structure, because Dirichlet mixture models with large number of components are inconsistent with respect to the true number of clusters [11]. Fig. 14 shows that both the number of larger clusters and the number of all clusters is stable between the different settings, when the model is saturated (with 18 or more global clusters) and provided that the corresponding chain reached convergence. The only exception is the solution with 45 global clusters where the algorithm identified an alternative clustering solution, with lower likelihood. In this case the probabilistic algorithm remained in a local optimum. We provide additional analysis of convergence of the presented algorithm in S1 Appendix.

Even though the number of estimated clusters changes with parameter settings, the cluster structures might still be very similar. In order to explore these similarities, the ARI between pairs of clusterings of different sizes is displayed graphically in Fig. 12. As the area of high ARI values (that is, similarity of clusterings) in the upper right corner indicates, the solution with 18 global clusters is a part of a larger group of solutions with similar cluster assignments. The result with 18 global clusters corresponds to the maximal log likelihood also corresponds to the smallest number of global clusters for which the results stabilise and then remain consistent for larger numbers of global clusters. The result with 45 global clusters is again an outlier.

**Figure 12.**
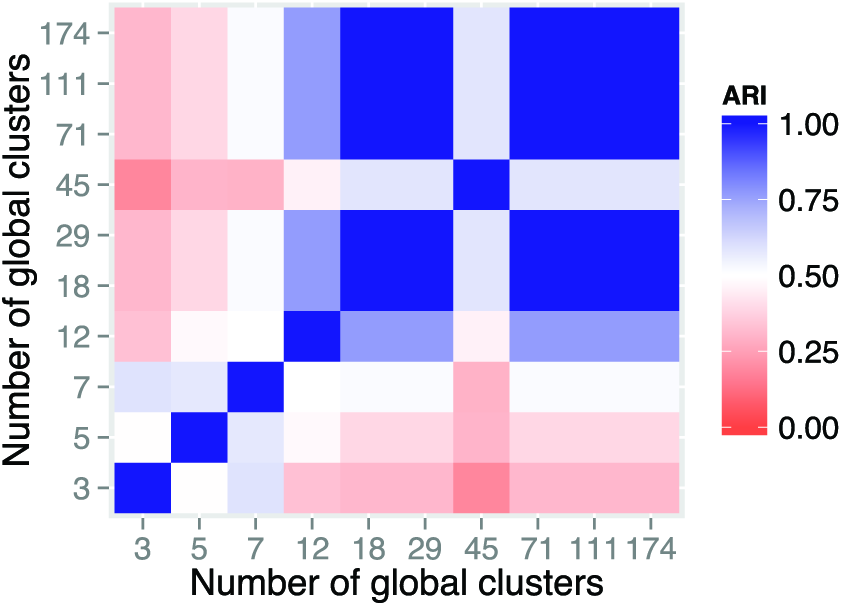
Consistency between global clustering results for different number of global clusters with 3 context-specific clusters, as measured by the ARI.

Fig. 13 shows the pairwise ARI values for local cluster assignments. The local assignments become stable at various sizes of the model. Generally, the larger number of possible global clusters give the model more flexibility to model cluster structures in individual datasets, as opposed to modelling a common global structure. Again, 18 global clusters represent the parameter setting where all the context-specific clusters converge to similar cluster assignments.

**Figure 13.**
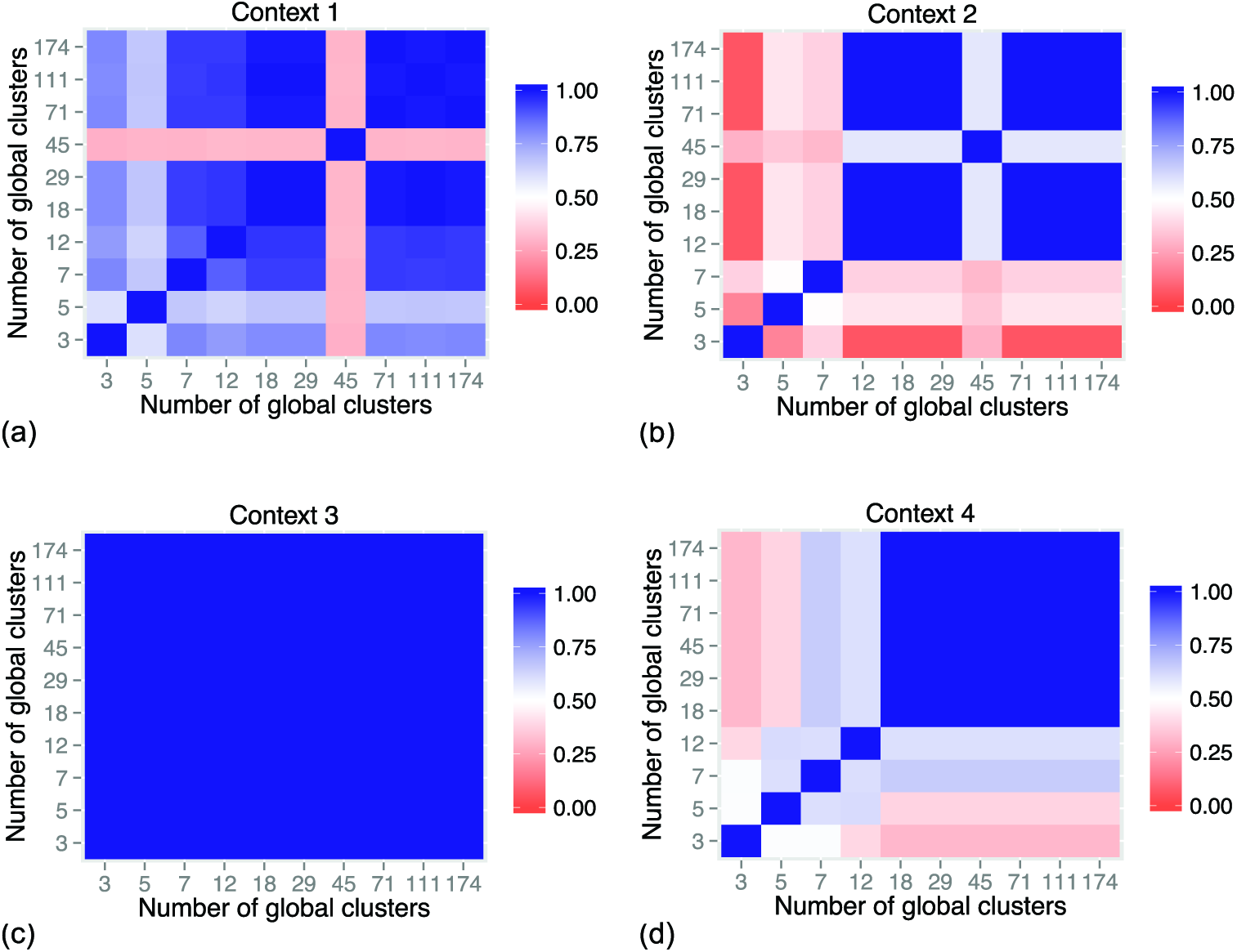
Consistency between local clustering results for different number of global clusters with 3 context-specific clusters, as measured by the ARI. The ARI values show several local optima. (a) Gene expression context. (b) DNA methylation context. (c) miRNA context. (d) RPPA context.

**Figure 14.**
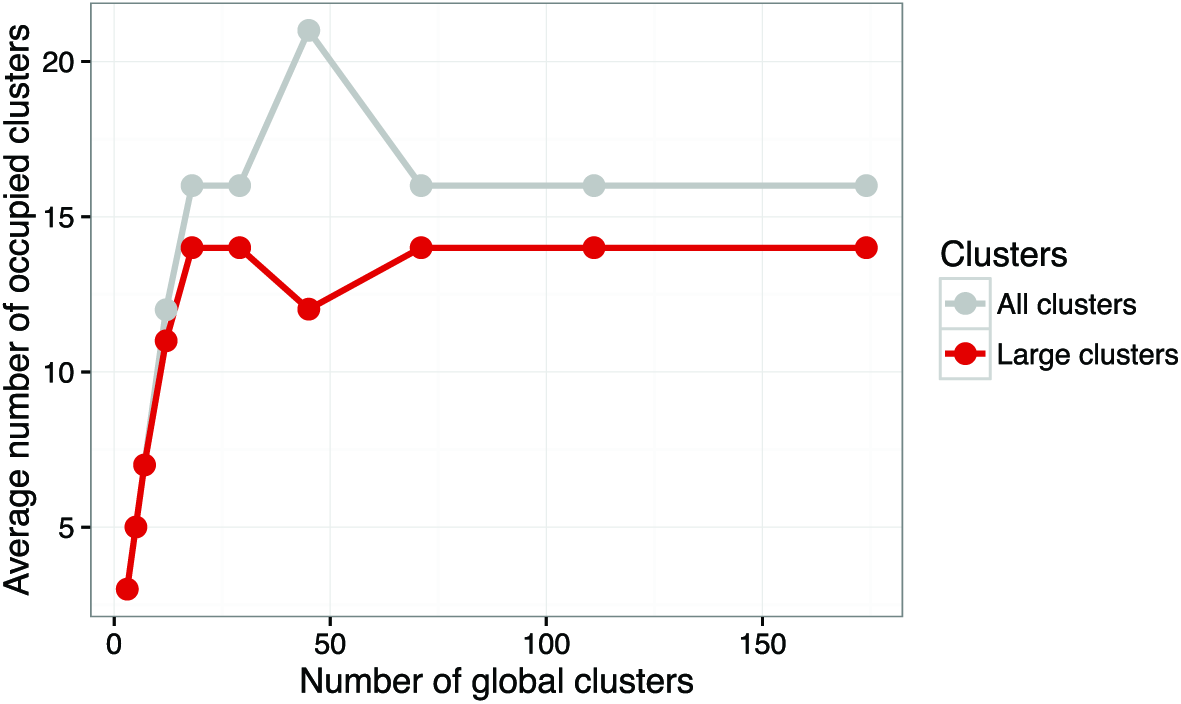
Average number of occupied clusters across different numbers of global clusters. The number of clusters is the average of the posterior number of global clusters that have any samples assigned to them across the MCMC iterations. The figure shows both the total number of occupied clusters and the number of clusters that have more than 5 samples assigned to them.

We also look at the effect of setting different numbers of local clusters. We look in detail at 3, 4 and 5 local clusters in every local context-specific dataset. Fig. 15 shows the agreement between the results as measured by the ARI. All the cluster assignments are relatively consistent given different model sizes with high pairwise ARI values.

**Figure 15.**
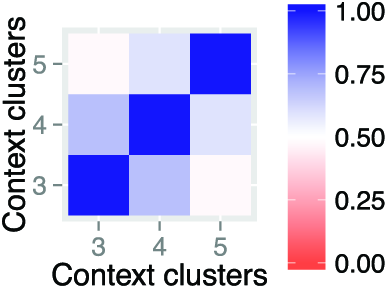
Consistency between global clustering results for different number of local context-specific clusters, as measured by the ARI. The compared models were trained with 18 global clusters and 3 to 5 context-specific clusters.

In general, the number of clusters tends to saturate for larger sizes of the model. Because the model asymptotically approaches a Dirichlet process when the number of global/local clusters is large, the model automatically infers the number of clusters that is needed to represent the data (see the Methods section for details). However, smaller sizes of the model are more computationally efficient in real-world scenarios. To infer a reliable clustering of real-world data, it is necessary to explore several different settings of the model's parameters, where stability of the clustering can serve as an indicator of clustering tendencies in the dataset [13].

To assist with selecting the number of clusters, the package *clusternomics* also provides the Deviance Information Criterion (DIC, [14]). DIC is a criterion for model selection for Bayesian models, combining posterior likelihood with penalty for model complexity. Number of clusters should be chosen so that the value of DIC is minimized. Fig. 16 shows the DIC values for the number of global clusters for the breast cancer dataset. Based on this measure, the optimal number of clusters is 18, which is also the value where the cluster results become stable.

**Figure 16.**
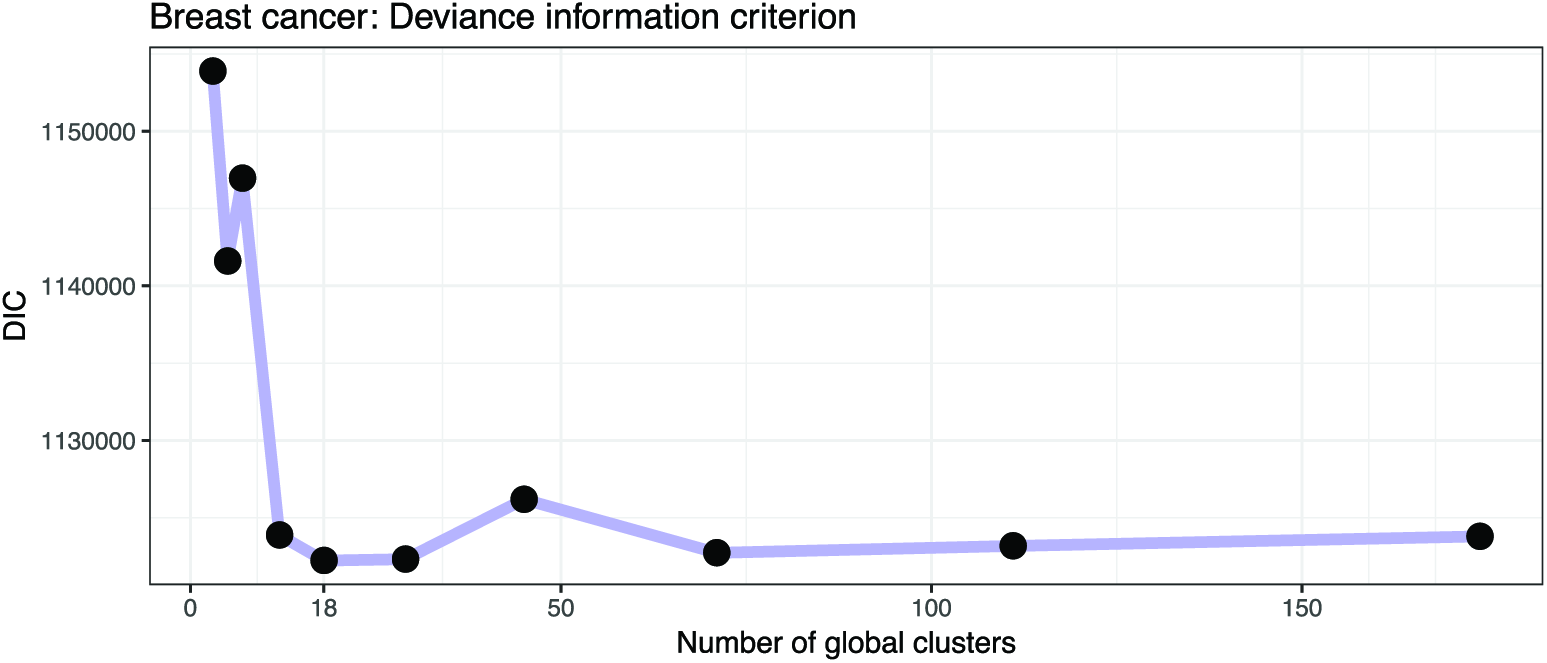
Deviance information criterion (DIC) as a method for selecting number of clusters. The plot shows the DIC for a range of numbers of global clusters when the number of local clusters is set to three. The DIC is minimized for 18 global clusters.

### Case Study: Prognostic Clusters in Lung and Kidney Cancer

To evaluate the general utility of the model, we also examined two additional smaller cancer datasets from The Cancer Genome Atlas repository:

- Lung cancer samples from 106 patients with 3 contexts: gene expression (12,042 genes), DNA methylation (23,074 loci) and miRNA expression (352 miRNAs)
- Kidney cancer samples from 122 patients with 3 contexts: gene expression (17,899 genes), DNA methylation (24,960 loci) and miRNA expression (329 miRNAs)

The datasets were normalised to zero mean and unit variance for every feature, but the total size of the data was not reduced. For both datasets we again fit models with 3 to 5 local clusters in each context and varying numbers of global clusters.

In this case, the size of the problem in terms of number of genes and DNA methylation loci is problematic for the iCluster algorithm. For larger problems with more features it becomes increasingly memory intensive (at least 60 GB of memory were required to run iCluster on the lung dataset, and the algorithm did not terminate in a reasonable period of time).

Fig. 17 shows the summary of the results on the lung cancer dataset, using the clinical survival information. The differences in survival prospects in the clusters are statistically significant for all clusterings of this dataset. The consistency of results reveals that there are three versions of stable cluster assignments across the models of different sizes. Fig. 19 shows similar results for the kidney cancer dataset. Here, the survival *p*-values drop below the significance threshold when the cluster assignments become more stable and consistent. In both cancer datasets, the Clusternomics algorithm identified clinically relevant clusters.

**Figure 17.**
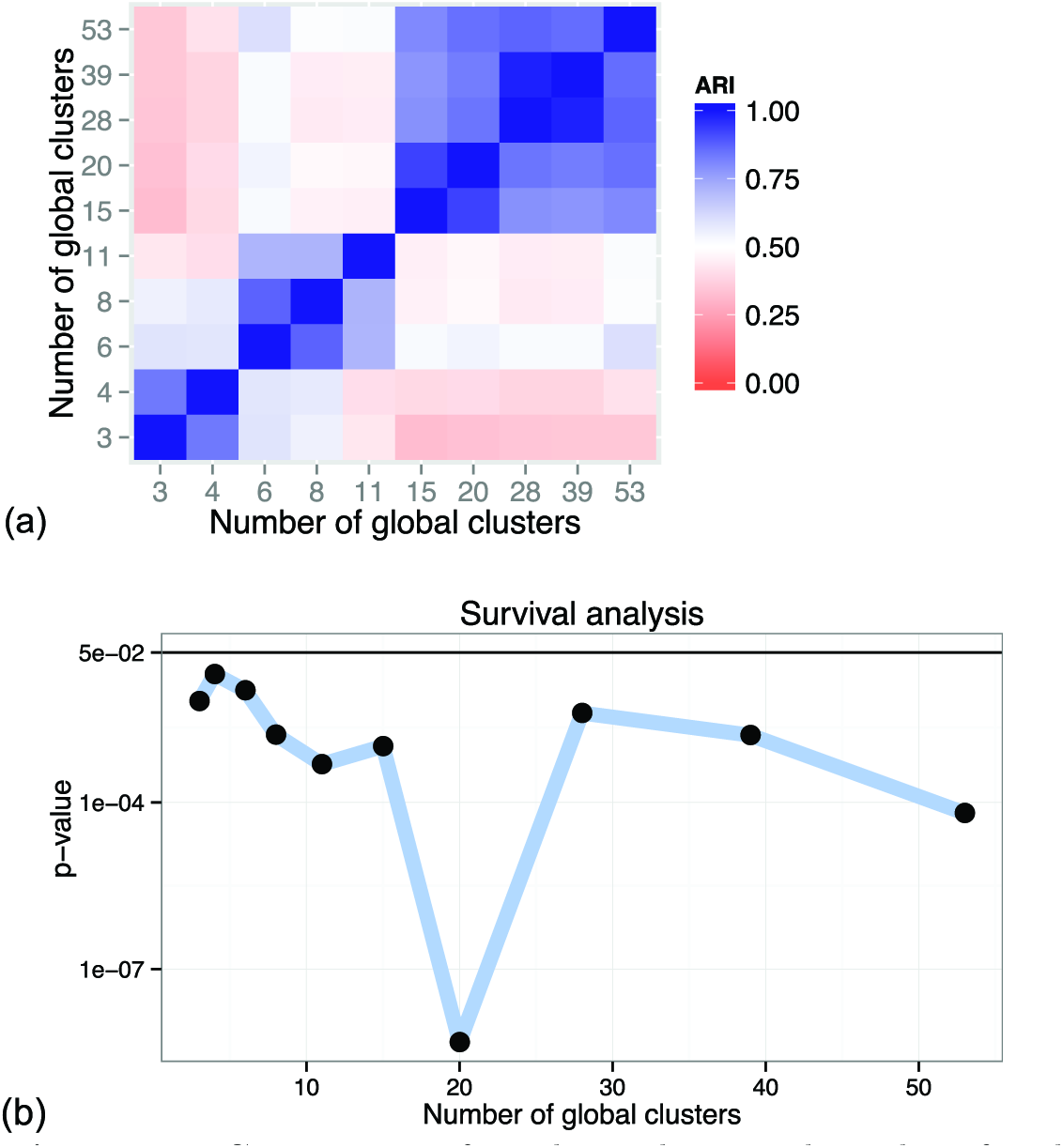
Consistency of results and survival *p*-values for clusters identified in the Lung cancer dataset, with a range of numbers of global clusters and 3 local clusters. (a) Consistency of results with respect to the ARI between different settings of numbers of global clusters. (b) *p*-values corresponding to the different numbers of global clusters.

**Figure 18.**
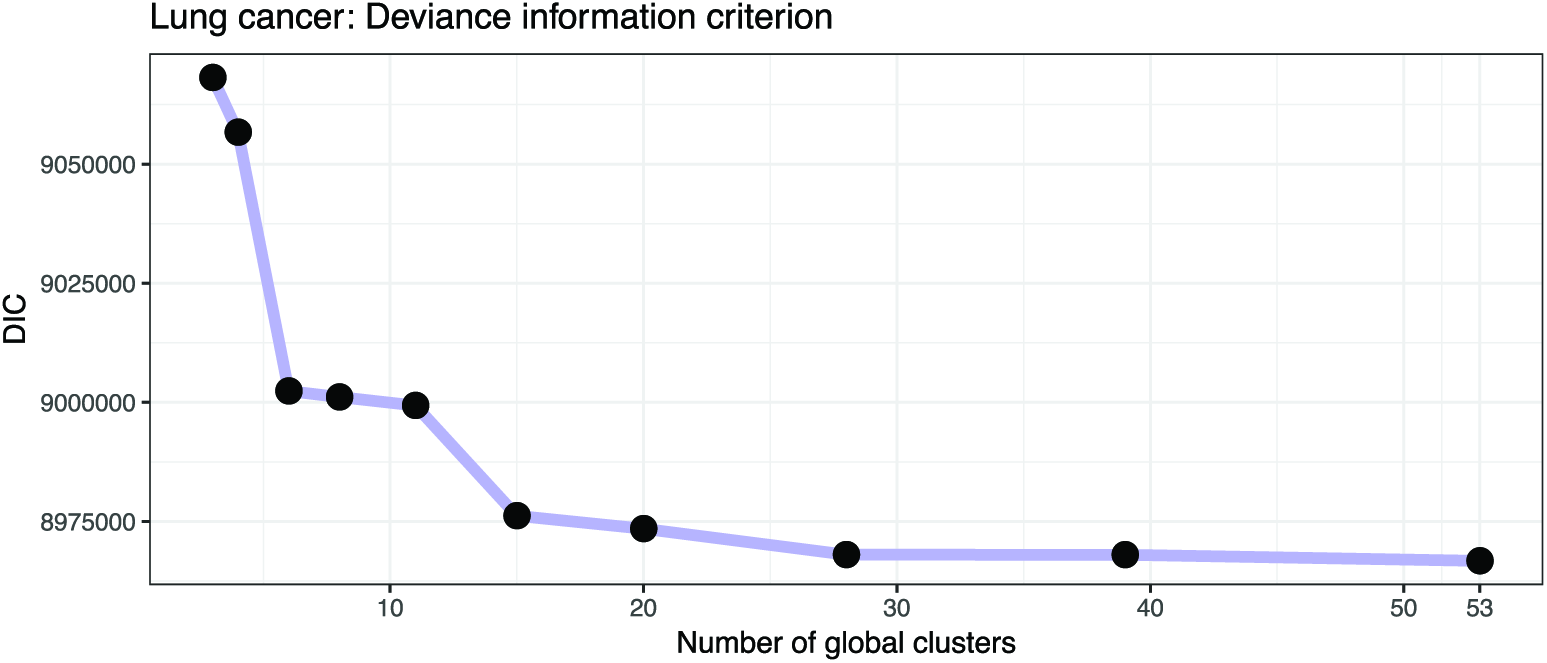
Deviance information criterion (DIC) for selecting number of clusters in the Lung cancer dataset. The plot shows the DIC for a range of numbers of global clusters when the number of local clusters is set to three. The DIC is minimized for 53 global clusters.

**Figure 19.**
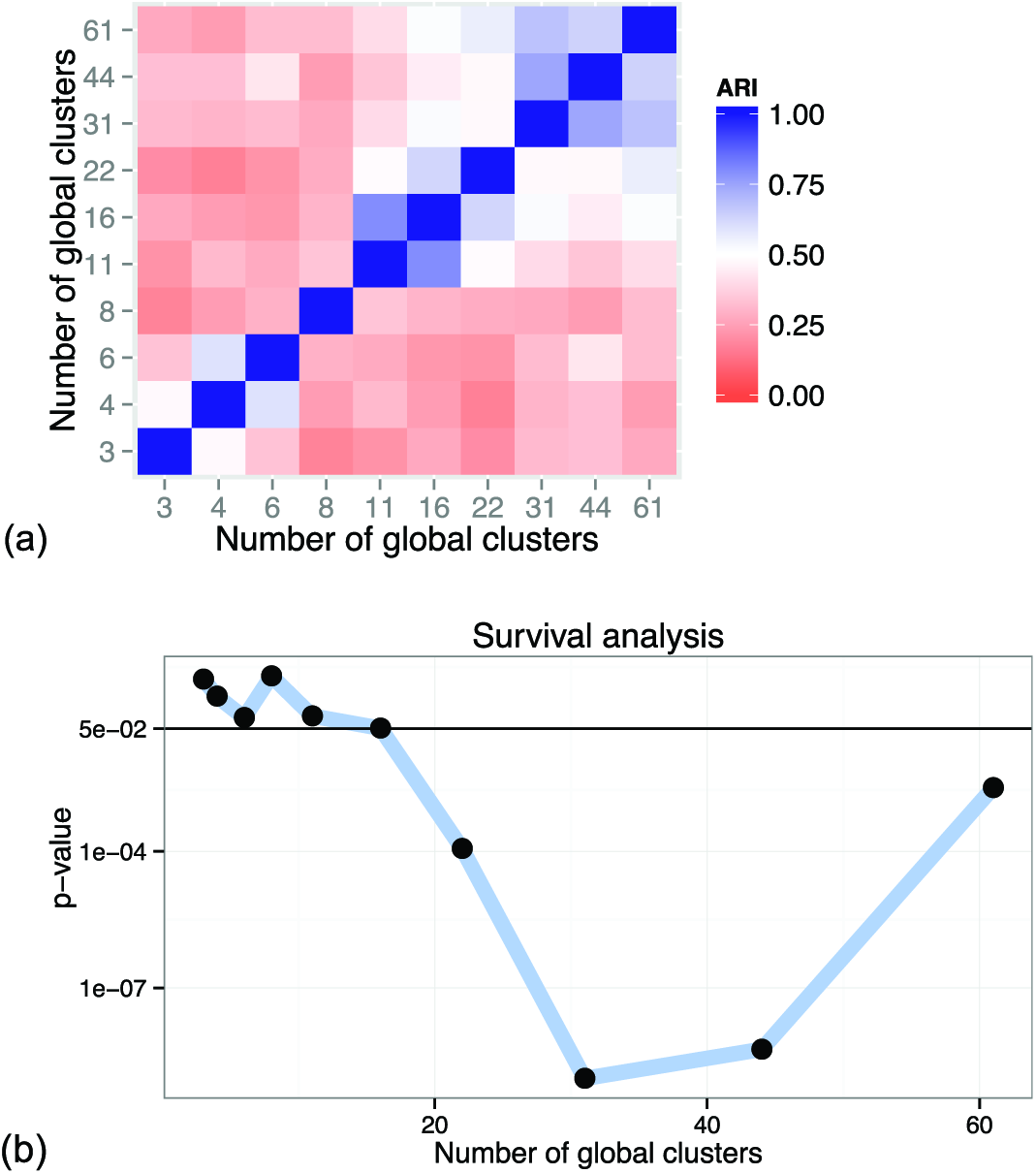
Consistency of results and survival *p*-values for clusters identified in the Kidney cancer dataset, with a range of numbers of global clusters and 3 local clusters. (a) Consistency of results with respect to the ARI between different settings of numbers of global clusters. (b) *p*-values corresponding to the different numbers of global clusters.

## Methods

### The context-dependent clustering model

The context-dependent clustering model explicitly represents both the local clusters within each dataset (local context), and the global structure that emerges when looking at the combination of cluster assignments across the individual datasets.

When we consider a local structure within several datasets, each dataset has its own context-specific set of clusters. When we look at the combination of the context-specific clusters, we get a combined structure which defines clusters on the *global* level while keeping information about cluster membership on the *local* level. For example, if one context (dataset) contains two clusters of samples labelled 1 and 2, and second dataset contains three clusters, labelled *A, B* and *C*, we get a combined structure with six potential global clusters where each cluster corresponds a combination of assignments on the local level:

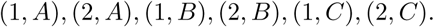

Figure 21 provides a schematic illustrat ion of the concepts. Using this formulation, we can model groups of data that are joined in one context and separated in another context, because they correspond to different global clusters.

**Figure 20.**
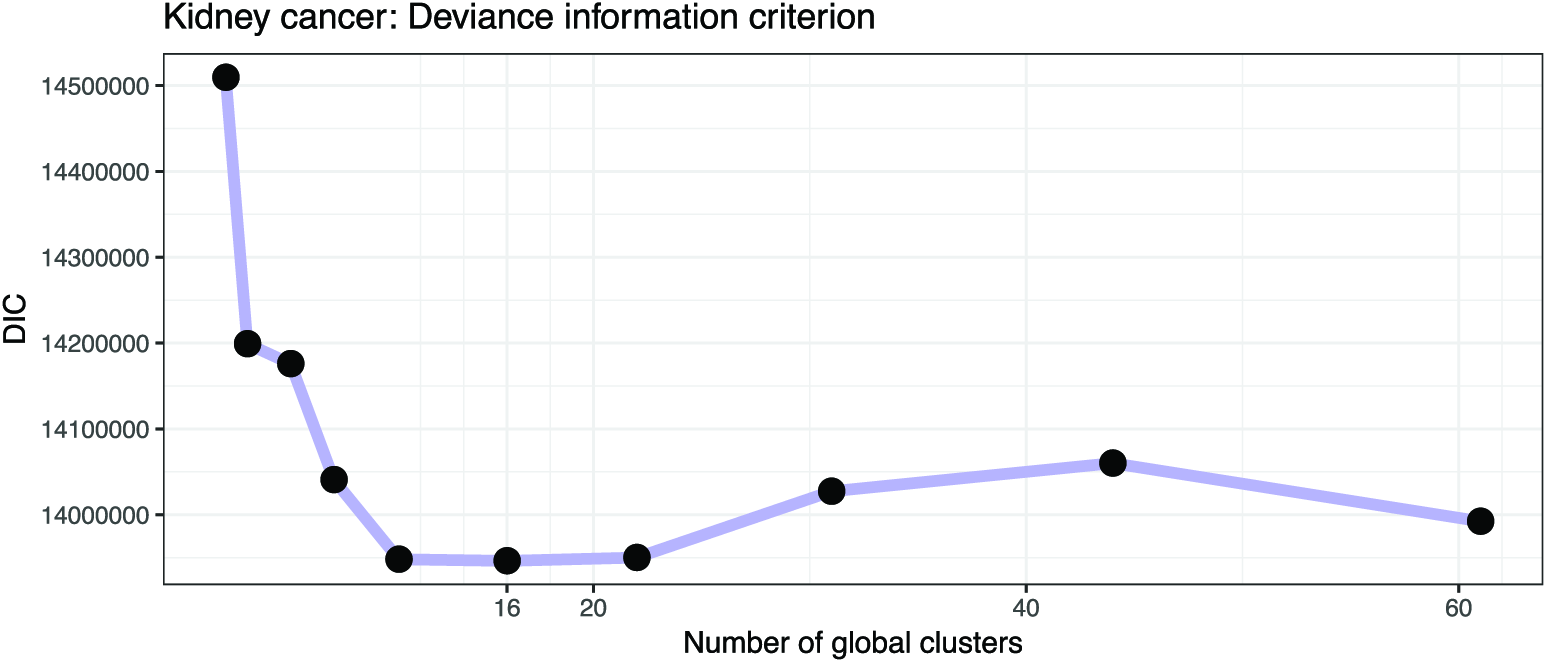
Deviance information criterion (DIC) as a method for selecting number of clusters in the Kidney cancer dataset. The plot shows the DIC for a range of numbers of global clusters when the number of local clusters is set to three. The DIC is minimized for 16 global clusters.

**Figure 21.**
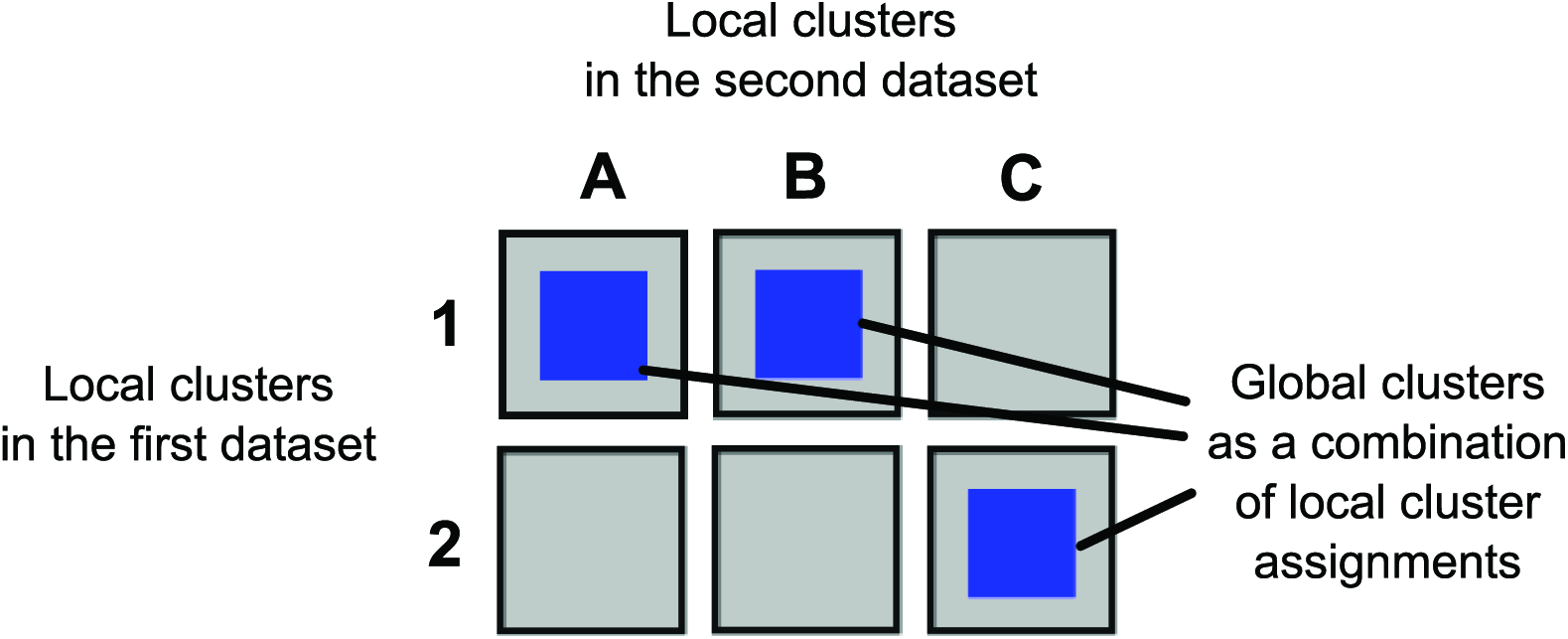
Illustration of the concepts of *global* and *local* clusters. The first dataset contains two clusters 1 and 2, the second dataset contains three clusters, *A, B* and *C*. The combined structure contains six potential global clusters that correspond to combinations of assignments on the local context level.

We use this intuition to develop the context-dependent clustering model, which is based on a Bayesian probabilistic clustering. The algorithm is based on Dirichlet mixture models [15] and their infinite generalisation, the Dirichlet process mixture model [16, 17].

The probabilistic model accounts both for probabilities of individual samples belonging to a specific cluster on the local level, and for the global probabilities of samples belonging to a global cluster. The global clusters are modelled as a combination of local clusters, and bound together by a Bayesian hierarchical model. This assures that a local cluster assignment changes the posterior probabilities of corresponding global cluster assignments.

Continuing with the two dataset example, when a sample is assigned to cluster 1 in the first dataset, it increases the posterior probability of the cluster within the first dataset, but it also increases the probability of global clusters (1, *A*), (1, *B*) and (1, *C*). This dependence is defined by the Bayesian hierarchical model, and encodes the objectives stated in the Introduction.

### Model description

In this section we introduce two alternative Bayesian probabilistic models for context-specific clustering. The first model looks explicitly at all possible combinations of local clusters which define the global combinatorial clusters. The second formulation of context-dependent clustering only models a restricted set of combinations of local clusters, that are data-driven.

Both models are asymptotically equivalent (for details see 81 Appendix). The first formulation of the model provides an intuition that leads to the second formulation of the model. Because the second formulation uses only a smaller number of local clusters, it is more computationally efficient than the first formulation, and it was therefore used to compute the results presented in the Results section.

We introduce the following notation: *X_n_*, *n* = 1, …, *N*, are data items where *X_n_* is composed of a set of observed values coming from contexts *c* = 1, …, *C*:

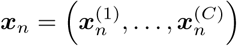

For example, for *C* = 2, *x_n_* may represent a tumour sample from patient *n* with gene expression values *x_n_*^(1)^ and DNA copy number states *x_n_*^(2)^.

#### Basic Dirichlet mixture model

The basis of the integrative hierarchical model is standard Bayesian model-based clustering with Dirichlet prior [15]. This basic model is used to cluster data within each context *c*.

This clustering model has been previously applied to gene expression studies, see for example Medvedovic et al. 2002 [18]. Lock and Dunson 2013 [7] used the same model as the basis of their integrative BCC model.

*K*^(*c*)^ is a fixed parameter for the maximum number of clusters in dataset *c* and *z_n_*^(*c*)^ is an indicator variable which defines the cluster assignment of sample *x_n_*^(*c*)^, 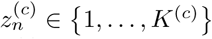. The probability of a sample belonging to a cluster *k* in context *c* is 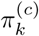,

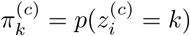

Values 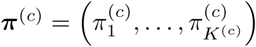 define the weights of each mixture component and follow a Dirichlet distribution with concentration parameter *α*_0_

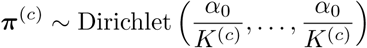

We define a finite mixture distribution for samples 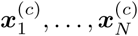 as

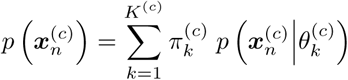

where 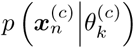 defines a probability distribution for sample 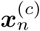 under mixture component 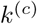 with parameters 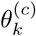.

The distributions in the mixture model for each context care summarised as follows:

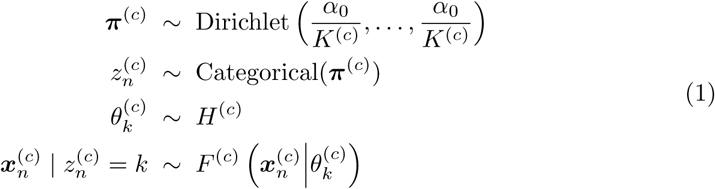

where *H*^(*c*)^ is some base prior distribution for parameters of each mixture component; *F*^(*c*)^ is a probability distribution for samples given parameters 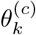. Note that the parameters *θ*^(*c*)^ and data distribution *F*^(*c*)^ depend on the context *c* and therefore can be modelled differently for different contexts. In general, we can also use different concentration parameters *α_c_* for each context.

### First model formulation: fully combinatorial model

Using the basic model presented in the preceding section we can now construct a composite model for integrative clustering, which we call Context-Dependent Clustering (CDC). To keep the notation simple, we first present the model for only two contexts, *c* ∈ {1, 2}. We start by specifying two mixture distributions as defined in the previous section, one for each context. Each context has its own mixture weights *π***^(1)^** and *π***^(2)^**, with symmetric Dirichlet priors:

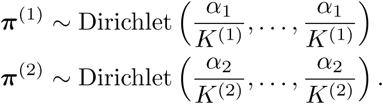

These two mixture distributions form the basis of *local* clustering within each context. We link the two distributions together using a third mixture distribution, also with a Dirichlet prior over the mixture weights *ρ*. This represents the *global* mixture distribution, defined over the outer product of *π*^(1)^ and *π*^(2)^:

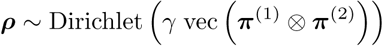

Here the outer product of *π*^(1)^ and *π*^(2)^ is

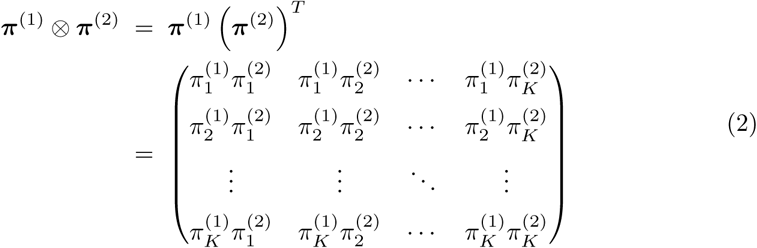

The vec operation takes a matrix and stacks its columns on top of each other to form one column vector.

We use the outer product matrix in a vectorised form as the basis for the (non-symmetric) concentration parameters of the Dirichlet distribution over *global* mixture weights *ρ*. Each element of *ρ* corresponds to a specific pair of local cluster probabilities *π_i_*^(1)^ and *π_j_*^(2)^. This effectively creates a mixture model over all possible combinations of cluster assignments on the level of individual contexts. The prior probability of a data item being simultaneously assigned into cluster *k* in the first context and cluster *l* in the second context corresponds to the element *s* in the ***ρ*** vector which originated from the row *k* and column *l* in the outer product matrix (2):

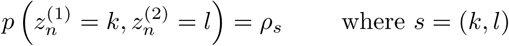

We also define a composite cluster indicator variable *z* for each sample that represents a pair of *z*^(*c*)^ values, one for each context *c*:

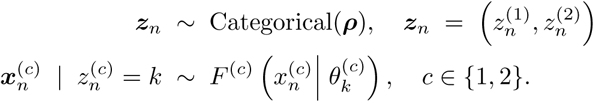

By fitting this model to a dataset, we obtain both local clusters on the level of the original datasets (contexts) and the global composite clusters that represent different combinations of individual local cluster assignments. Local clusters for context *c* can be obtained by taking a projection of the *z* indicator variable onto the context *c*.

Going back to the HER2 oncogene example in the Introduction, the model explicitly represents the local clusters with respect to DNA copy number changes and mRNA expression, while forming two global clusters with respect to the overall behaviour.

The model satisfies the objectives presented in the Introduction. In the posterior, assignments into a context-specific cluster affect the posterior distribution of *π*^(*c*)^, which in turn affects the probabilities of all global combinatorial clusters through the hierarchical model. For example if we look at the outer product matrix (2) which is a part of the prior for global mixture weights ***ρ***, change in the value of *π_k_*^(1)^ changes values of the whole *k*-th row in the matrix.

The model also represents different degrees of dependence by allowing any combination of cluster assignments across contexts. When there is a single common cluster structure across the two contexts, the occupied clusters will be concentrated along the diagonal of the probability matrix (2).

For *C* > 2, the outer product of 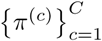 generalises into a tensor product. Each element of *ρ* represents the probability of a *C*-tuple of cluster assignments, i.e. a specific combination of cluster assignments in specific contexts. To summarise the model, we give a general formulation for *C* contexts.

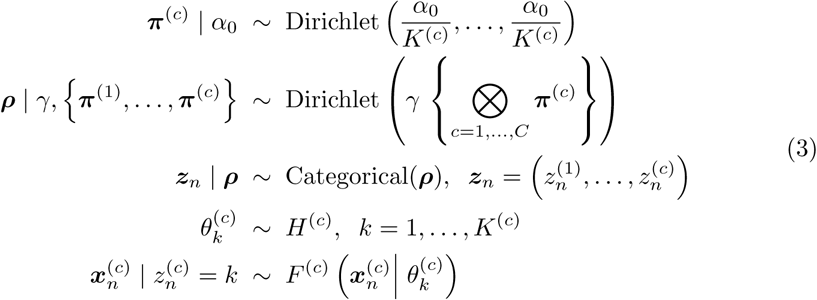

Fig. 22 shows the graphical model for the full context-dependent clustering model. In general, the number of clusters *K*^(*c*)^ can be different for each context *c*. Given that the number of clusters in each dataset is 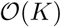, the total length of ***ρ*** parameter vector is 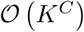 because the model represents all possible combinations of cluster assignments. This yields a very large number of potential combinatorial clusters. However, only a small number of clusters is actually represented in the data and many of the clusters remain empty as shown by Rousseau et al. [19]. Also, by using small values for the concentration parameter *γ* < 1, we encourage the data to be concentrated in only a small number of global mixture components.

**Figure 22.**
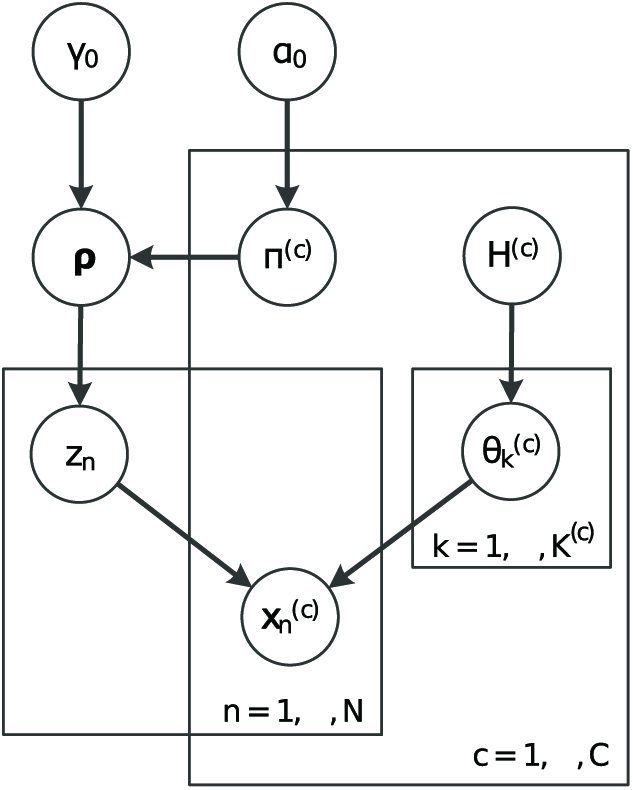
Graphical model representation of the fully combinatorial context-dependent clustering model.

This model explicitly represents all the possible combinations of cluster assignments, which may not be desirable in real-world applications, where the model may use some of the structure to capture technical noise present in the data. Also, although the number of clusters that are occupied is smaller in the posterior, the model size grows exponentially with the number of contexts. In the next section, we look at an alternative model that circumvents this limitation.

### Second model formulation: decoupled combinatorial model

To avoid the large number of potential cluster combinations in the previous model, we decouple the number of context-specific clusters *K*^(*c*)^, *c* = 1, …, *C*, and the number of global clusters. First a mixture distribution over *S* global clusters is defined similarly to the finite Dirichlet mixture model [15].

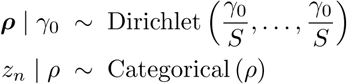

where ***ρ*** are the mixture weights and *z_n_* are standard cluster assignment indicator variables. We combine context-specific clusters with the global clusters using the following method: we specify context mixture distributions and a set of assignment variables *k_s_*^(*c*)^ that associate global cluster *s* = 1, …, *S* with context-specific clusters:

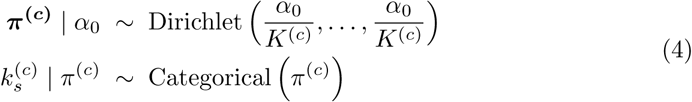

In this formulation, 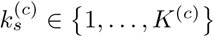 assigns the *s*-th global cluster to a specific local cluster in context *c*. One could say that 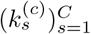 are the coordinates of the global cluster in terms of the local cluster identifiers. The variables *z_n_* then assign samples to global clusters. This way, data are represented by a mixture of context-specific cluster combinations. In the previous model (3) the mapping of global clusters to local clusters was implicit, because each combination of context clusters mapped to a unique global cluster. In this model, the mapping is probabilistic and forms a part of the model.

Note that compared to the previous model, we also have to specify the number of potential global clusters *S* which is no longer determined by the number of clusters within each context. This circumvents the problem of large dimensionality of the space of potential cluster combinations across contexts.

To summarise, the complete model can be written as

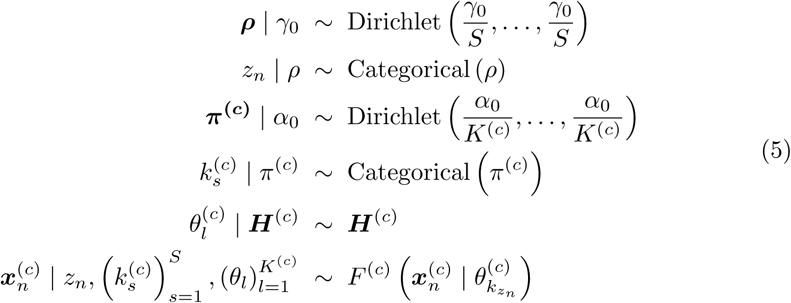

Fig. 23 shows the graphical representation of this model.

**Figure 23.**
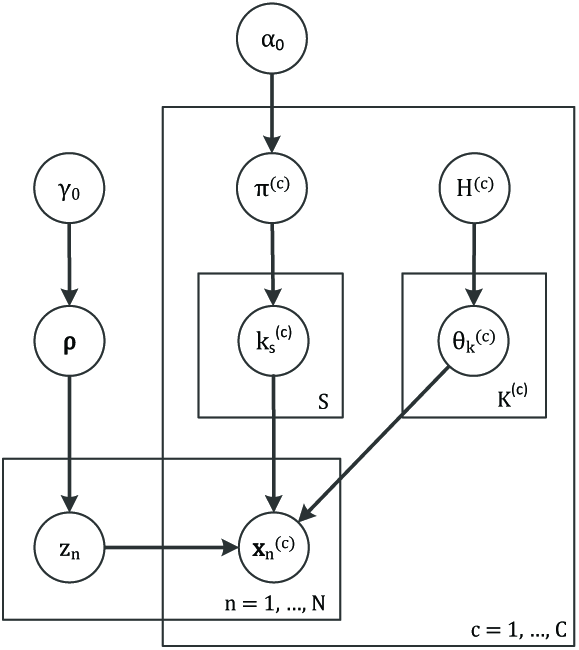
Graphical model representation of the decoupled context-dependent integrative clustering model.

### Inference in the model and implementation

We derived Gibbs sampling inference algorithms for both formulations of the context-dependent clustering model, details and the inference equations can be found in S1 Appendix.

For the fully combinatorial version of the model, computational complexity of each iteration of the Gibbs sampling algorithm is 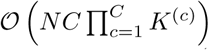, additionally multiplied by the complexity of evaluating the data likelihood *F*^(*c*)^ for each context. We also derived approximate variational inference updates for this version of the model, which achieves faster convergence than Gibbs sampling. Details on variational inference in the model are also available in S1 Appendix.

For the decoupled formulation of the model, the computational complexity of each Gibbs sampling iteration is 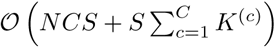, again additionally increased by the complexity of evaluating data likelihood *F*^(*c*)^. Compared to the fully combinatorial model, complexity of this algorithm is lower for 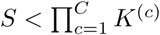 because of the decoupled representation.

Implementation of the decoupled version of the context-dependent clustering model, which was used to produce the results presented in the Results section, is available as the R package `clusternomics` from CRAN.

## Discussion

To summarize, in this paper we proposed a clustering algorithm for integrative analysis of heterogeneous datasets. We described the probabilistic model behind the algorithm, which is closely related to the hierarchical Dirichlet process [20]. The proposed context-dependent clustering algorithm models both the local structure within each dataset, and the global structure which arises from combinations of dataset-specific clusters. This form of model enables modelling of heterogeneous related datasets that do not share the same structure.

We described two representations of the model which are equivalent in their limit. The first full model makes the assumptions behind the model explicit and represents all possible combinations of context-specific cluster assignments. Given the number of clusters in each context *K*^(*c*)^ the model is a mixture model over all possible combinations of cluster assignments in individual contexts.

The second type of representation is the decoupled CDC model which allows us to specify the number of global clusters *S* that are identified in the data separately from the number of context-dependent clusters *K*^(*c*)^. For a large number of global clusters *S* the model is equivalent to the full model. However, the number of global clusters allows us to additionally tune the resulting cluster structure. For smaller numbers of clusters *S*, the global and local cluster structures are forced to be more similar and the algorithm enforces a common cluster structure across all datasets. For larger numbers of global clusters, the model has more flexibility to model the local structure within each dataset.

We evaluated the proposed model both on simulated data and on a set of real-world cancer datasets. The simulated data revealed that other algorithms for integrative clustering do not model situations where there is a varying degree of dependence of cluster structures across multiple datasets.

We also evaluated the proposed context-dependent clustering model on the breast cancer dataset which includes four different contexts, and additionally on two datasets studying lung and kidney cancer. The model successfully identified clinically meaningful clusters as measured by the survival probabilities for each global cluster. We evaluated the decoupled clustering model over a number of possible global clusters. Generally, the best clustering results are obtained when the cluster structure stabilises as measured by the ARI. Senbabaoglu et al. [13] note that in real-world datasets there may be many different numbers of clusters that are equally highly plausible. The comparison of clustering results based on their agreement identifies the model sizes that lead to similar sets of cluster assignments. Each group then corresponds to an alternative interpretation of the data.

Overall, the model uses different assumptions about the cluster structure than other currently used integrative clustering algorithms. By dropping the assumption of a single common cluster structure, the model identifies both the local structure of individual datasets, and a global structure that combines the local clusters.

## Supporting Information

### S1 Appendix

#### Model and implementation details

Details on the model, implementation, algorithm setting and MCMC convergence.

## Acknowledgments

The authors would like to thank David L. Wild and Richard E. Turner for helpful comments.

